# Woody plant taxonomic, functional, and phylogenetic diversity decrease along elevational gradients in Andean tropical montane forests: environmental filtering and arrival of temperate taxa

**DOI:** 10.1101/2023.08.05.551864

**Authors:** Guillermo Bañares-de-Dios, Manuel J. Macía, Gabriel Arellano, Íñigo Granzow-de la Cerda, Julia Vega-Álvarez, Itziar Arnelas, Carlos I. Espinosa, Norma Salinas, Luis Cayuela

## Abstract

**Aim:** Mountains are paramount for exploring biodiversity patterns and their causes due to the rich mosaic of topographies and climates encompassed over short geographical distances. Biodiversity changes along elevational gradients have traditionally been explored in terms of taxonomic diversity, but other aspects must be considered. For first time, we simultaneously assessed elevational trends in the taxonomic, functional, and phylogenetic diversity of woody plants in Andean tropical montane forests (TMFs) and explored their underlying ecological and evolutionary causing processes.

**Location:** Tropical Andes

**Time period:** 2011/2012 and 2017/2019

Tropical Andes

**Major Taxa:** Woody plants

**Methods:** We investigated taxonomic, functional, and phylogenetic diversity along four transects (traversing *ca*. 2,200 m altitudinal gradients) encompassing 114 0.1 ha plots across a broad latitudinal range (*ca*. 10°). We used Hill numbers to quantify differences in the abundance-based diversity of 37,869 woody plant individuals with DBH ≥ 2.5 cm.

**Results:** Taxonomic, functional, and phylogenetic diversity decreased as elevation increased. The decrease was less pronounced for Hill numbers of higher orders. The only exception was a slight increase in phylogenetic diversity when more weight was given to dominant species. These results were consistent between transects.

**Main conclusions:** The decrease in taxonomic and functional diversity with elevation might be due to an environmental filtering process where the increasingly harsher conditions towards highlands exclude species and functional strategies. Besides, the differences in the steepness of the decrease between Hill orders suggest that rare species contribute disproportionately to functional diversity. The shifting elevational trend in the phylogenetic diversity between Hill orders indicates a greater than previously considered influence in central tropical Andean highlands of species originated in lowlands with strong niche conservatism relative to distantly related temperate lineages. This could be explained by a decreasing presence and abundance of extratropical taxa towards the central Andes relative to northern or southern Andes.

**BIOSKETCH:** Guillermo Bañares-de-Dios is a plant ecologist with interests in community assembly, biodiversity patterns, and global change. He completed his PhD in 2020 and belongs to “Grupo de Ecología Tropical”, an international network of researchers from different institutions with broad interests in tropical biology (http://www.grupoecologiatropical.com/?lang=en). Currently he works as Project Manager implementing the European Pollinator Monitoring Scheme in Spain.

## 1 | INTRODUCTION

Biodiversity distribution patterns and their causes have been central to natural history and ecology since more than two centuries (Caldas 1803) until the present (Sanmartín 2012). Mountains provide invaluable insight for developing a general theory on biological diversity to explain these patterns (Körner 2004, Sanders and Rahbek 2012). Compared to latitudinal gradients, elevational gradients encompass more climate regimes, greater topographic complexity, and wider environmental oscillations over shorter geographical distances (Antonelli et al. 2018, Rahbek et al. 2019a). Thus, mountains are exceptionally biodiverse, where they comprise only 25% of Earth’s terrestrial surface but harbour *ca.* 87% of terrestrial biodiversity, with a large proportion of endemicity (Rahbek et al. 2019b). In particular, Andean tropical montane forests (TMFs) host one-sixth of all plant species, where 44% are endemisms (Myers et al. 2000). Furthermore, they play key ecosystem services such as carbon storage or water supply (de la Cruz-Amo et al. 2020). Therefore, Andean TMFs are among the biodiversity hotspots with highest conservation priority (Mittermeier et al. 2005). Lastly, from a biogeographic perspective, Andean TMFs are optimal natural laboratories to provide rigorous tests of diversity trends along altitude and comparisons of these trends among sites (Lomolino 2001, Guo et al. 2013, Graham et al. 2014) because they allow to replicate large elevational ranges over a continental north-south corridor which has witnessed historic processes across millions of years (Antonelli et al. 2009, Malhi et al. 2010, Tito et al. 2020).

Biodiversity has traditionally been explored in terms of taxonomic diversity by treating all Linnaean binomials as ecologically equivalent and evolutionary independent. This approach cannot yield much information about the ultimate drivers of biodiversity distribution patterns (Pavoine and Bonsall 2011). A more accurate depiction of diversity should also consider functional and phylogenetic diversity (McGill et al. 2006). Functional diversity uses species functional traits to understand where species can thrive, how they interact, or how they contribute to ecosystem functioning (Cadotte et al. 2011). Phylogenetic diversity tracks the origins of lineages of taxa considering the historical and evolutionary processes that shaped their distributions and thereby provides insights to explore the key events in biodiversity distribution (Cavender-Bares et al. 2009). In the particular case of the Andes, two hypotheses have been proposed to explain the formation of biological communities: a) the establishment of Amazonian clades originated under warm conditions at tropical lowlands –typically exhibiting strong conservatism of ancestral narrow thermal tolerances– which, during the Pleistocene, adapted to the comparatively cooler temperatures of the newly raised highlands resulting from the Andean uplift (Latham and Ricklefs 1993, Kerkhoff et al. 2014); and b) the arrival of cold pre-adapted lineages –originated much earlier in temperate latitudes– via migration along the Andean high-elevation corridor (Jablonski et al. 2006). The former (a) is the cornerstone of the Tropical Niche Conservatism (TNC, Wiens and Donoghue 2004) hypothesis, while the latter (b) is the core of the Out of the Tropical Lowlands (OTL, Qian and Ricklefs 2016) hypothesis. Despite these three diversity facets are intrinsically related (*e.g.*, closely related taxa tend to share more traits [Moles et al. 2005]) their responses to the environment can differ (Devictor et al. 2010). However, few studies have simultaneously considered the three types of diversity, particularly for plants under changing environmental conditions, especially along these inherent to elevational gradients (but see Tanaka and Sato 2015, Chun and Lee 2017).

Previous studies on biodiversity patterns along elevational gradients in different biomes and distinct taxonomic groups came to contrasting results regarding taxonomic diversity, but largely agreed in terms of functional and phylogenetic trends. Historically, the prevalent pattern for taxonomic diversity is a decrease with elevation, although some studies have described species richness peaks at medium elevations (Rahbek 1995, McCain 2005, Dani et al. 2023). The most common trend in functional diversity also comprises narrowing towards fewer functional strategies to cope with harsher environmental conditions as elevation increases (Swenson et al. 2012, de Bello et al. 2013). The elevational pattern for phylogenetic diversity is specific to each site′s biogeographic historical key events (Ricklefs and Schluter 1993). In the case of the Andes TNC predicts an elevational decrease in phylogenetic diversity whereas OTL predicts an increase.

In this study, we investigated plant diversity in Andean TMFs at four transects ranging elevational gradients from *ca.* 800 to 3,100 masl and spanning *ca.* 1,750 km in latitude across Ecuador, Peru, and Bolivia. We used census data from 114 plots (containing *ca.* 38,000 individuals and 2,000 species) and analysed them using Hill numbers to quantify different types of abundance-based diversity indices. We aimed to 1) assess the elevational trends in taxonomic, functional, and phylogenetic diversity, to 2) examine the relationships between different diversity facets along elevation and to 3) explore their underlying ecological and evolutionary causing processes. To the best of our knowledge, this is one of the most ambitious elevational studies of the three facets of plant diversity and a similar study has never been conducted before along the Andean range.

## 2 | MATERIALS AND METHODS

### 2.1 | Study area

We studied four transects ranging elevational gradients of Andean TMFs located in remote regions: Podocarpus National Park (Ecuador), Río Abiseo National Park (Peru), Madidi National Park (Bolivia), and Pilón-Lajas Biosphere Reserve (Bolivia) (Fig. 1). All four transects contained a similar elevational range (*ca.* 2,200 m), within which we defined three elevational belts: premontane (900–1,300 m), montane (1,800–2,250 m), and upper (2,700–3,100 m). Within each belt, we established nine to ten plots of 0.1 ha (50 × 20 m) as described by Arellano et al. (2016), with a total of 114 plots (Table 1 and Appendix 1, Table A1). Plots were located in mature and apparently undisturbed forests at distances of at least 300 m. In each plot we inventoried all woody stems (including trees, shrubs, lianas, hemiepiphytes, palms, and tree ferns, but excluding woody Poaceae) with a diameter at breast height (DBH) ≥ 2.5cm, with a total of 37,869 individuals (Table 1 and Appendix 1, Table A1).

**FIGURE 1.**
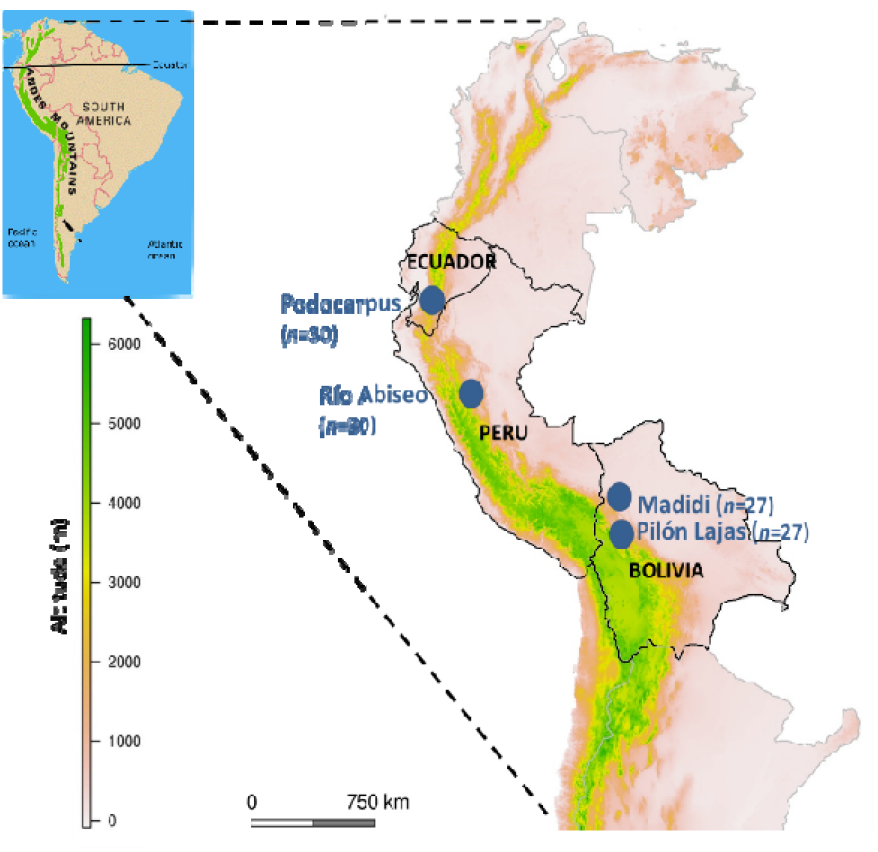
Locations (blue dots) of the four transects ranging elevational gradients in the tropical montane forests (TMFs) studied along the central Andes. Data from censuses of *n* = 114 TMF 0.1 ha plots located in remote regions: Podocarpus National Park (Ecuador), Río Abiseo National Park (Peru), Madidi National Park (Bolivia), and Pilón-Lajas Biosphere Reserve (Bolivia).

**TABLE 1.**
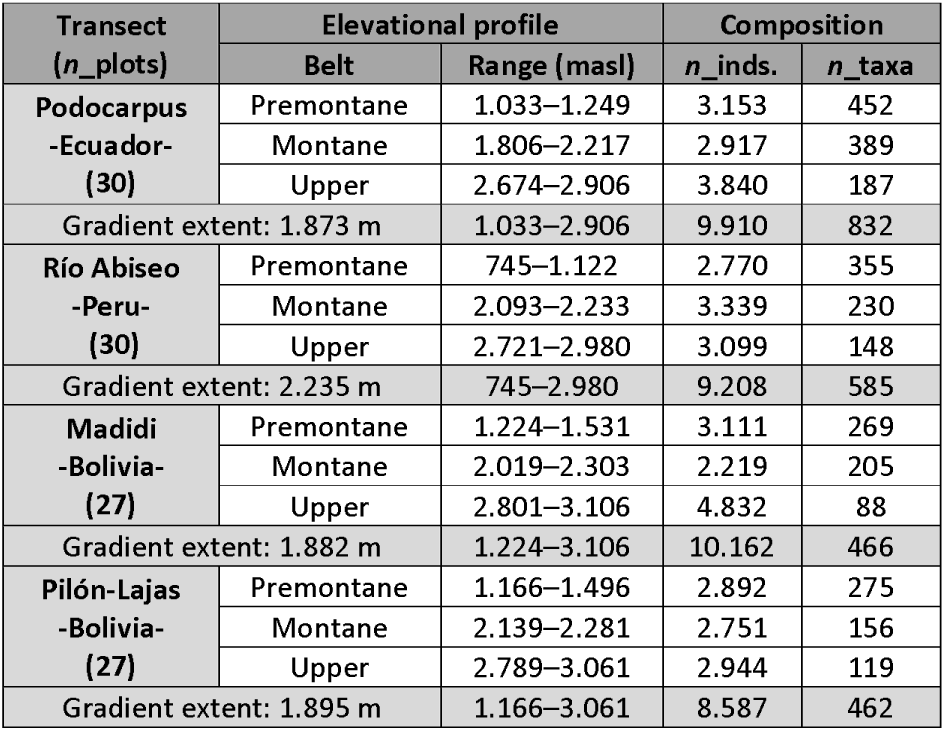
Summary of elevational profile and composition of the four transects ranging elevational gradients that encompassed 114 Andean TMFs plots. *n*_inds.: number of individual stems; *n*_taxa: number of taxa. Specific detail for each plot is available at Appendix 1, Table A1. All four transects contained a similar elevational range, within which we defined three elevational belts: premontane, montane, and upper.

Environmental characterization of the plots was conducted using bioclimatic variables from the CHELSA climatological dataset (Karger et al. 2017) (Appendix 1, Table A2). Mean annual temperature is the most representative variable for studies along elevational gradients since it represents the fundamental environmental change associated with elevation (Körner 2007). Mean annual temperature exhibits high correlation with elevation (*r*>0.98) in our study sites and the thermal ranges of transects varied slightly: 12°C in Podocarpus, 9°C in Río Abiseo, and 10°C in both Madidi and Pilón-Lajas. Alternative climatic variables related with precipitation are not as relevant for our study since 1) moist TMFs, characterized by persistent fog and high precipitation throughout most of the year, are not subjected to relevant seasonal water stress, and 2) at each transect, plots are located along the same river basin as to minimize spatial rainfall variability inherent to the complex topography of the Andes, thus leaving temperature as the key source of abiotic variation along our elevational gradients (Malhi et al. 2010). Small changes in mean annual temperature are expected to have a major influence on species distributions, especially in the tropics where species have narrow thermal tolerances (Janzen 1967).

**TABLE 2.**
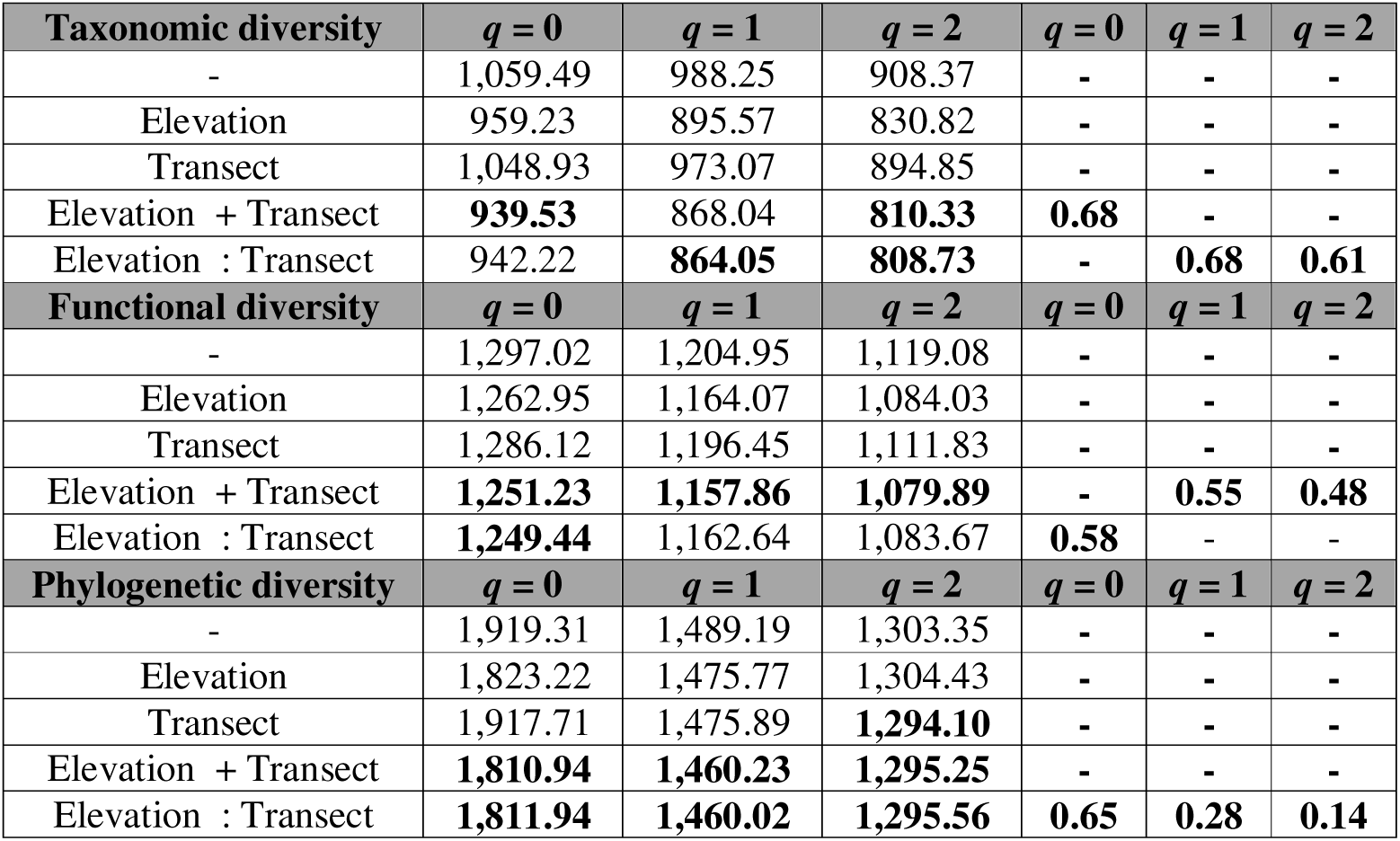
Corrected Akaike information criterion (AICc) values for generalized linear models (GLMs) of the taxonomic, functional, and phylogenetic diversity values calculated for Hill numbers of different order relative to distinct predictors for 114 plots of Andean TMFs. Values for the best fitted models are highlighted in bold. The explained deviance D2 was calculated only for the best and most complex model (if more than one model was the best, the differences in terms of deviance between the most and least complex would be minimal).

### 2.2 | Taxonomic and functional characterization

We collected voucher specimens in the field, which were later identified and deposited in local herbaria (HUTPL and LPB [Thiers, n.d.]). Taxonomic standardization was conducted using the ‘Taxonstand’ R package (Cayuela et al. 2012; v. 2.2). A summary of taxonomic composition for elevational belts can be found in Table 1 (specific details for each plot are available at Appendix 1, Table A1).

Functional characterization was only conducted for two transects: Podocarpus (Ecuador) and Río Abiseo (Peru). For each taxon, we measured the specific leaf area, leaf thickness, and branch wood density as described by Cornelissen et al. (2003). These functional traits have been widely used for assessing the key woody plant functional strategy axes (Chave et al., 2009; Wright et al., 2004). We calculated mean trait values for each taxon. On average for each plot, functional data for the three traits were available for 65 taxa (86%) in Podocarpus and 63 taxa (92%) in Río Abiseo.

### 2.3 | Phylogenetic tree generation

We generated a phylogenetic tree (Appendix 2) using the ‘V.PhyloMaker’ R package (Jin and Qian 2019; v. 0.1.0), which allowed us to prune our species list from the largest currently available dated mega-phylogeny for vascular plants.

For phylogenetic diversity calculations, we only considered taxa identified to the species or genus level, *i.e.*, 703 taxa (88%) in Podocarpus, 416 taxa (71%) in Río Abiseo, 453 taxa (97%) in Madidi, and 457 taxa (98%) in Pilón-Lajas. Among these taxa, in order to include those absent from the original mega-phylogeny in the tree, we joined them at the half-way point of the genus/family branch (for species/genera) using the parameter ‘scenarios’ = S3 in the ‘phylo.maker’ function.

### 2.4 | Diversity measurements

Attribute diversity comprises a unified framework for assessing biodiversity by integrating the three diversity facets: taxonomic, functional, and phylogenetic diversity (Appendix 1, Table A3) (Chao et al. 2014a). Each type of diversity is measured in different entities (species for taxonomic, species pairwise Euclidean distances for functional, and species tree branch length for phylogenetic), but all three can be transformed into directly comparable units by using generalized Hill numbers. Hill numbers quantify abundance-based taxonomic diversity using the parameter, *q*, which defines the sensitivity to species abundances (Chao et al. 2014b). For *q* = 0, all taxa are treated equally regardless of their abundance (*i.e.*, taxa richness), whereas larger *q* values give greater importance to the relative abundances of taxa (*i.e.*, more weight to abundant/dominant taxa and less weight to rare taxa). The attribute diversity framework extends the usage of Hill numbers to functional (Chiu and Chao 2014) and phylogenetic (Chao et al. 2010) diversity by weighting each entity based on either its relative functional distance compared with other taxa (functional diversity) or the relative length of the phylogenetic tree branch for that taxon (phylogenetic diversity). We calculated the attribute diversity indexes for Hill numbers of the orders *q* = 0, *q* = 1 and *q* = 2 at the plot level using the ‘renyi’ function of the ‘vegan’ R package (Oksanen et al., 2019; v. 2.5-6) for taxonomic diversity, the code from Chiu and Chao (2014) for functional diversity and the ‘ChaoPD’ function of the ‘entropart’ R package (Chao et al., 2010; v. 1.6-10) for phylogenetic diversity.

### 2.5 | Statistical analyses

First, to acknowledge the influence of sample size on diversity metrics we conducted Pearson′s correlations between number of trees per plot and the three facets of diversity (taxonomic, functional, and phylogenetic) for different Hill orders.

Next, we explored Pearson′s correlations among the three types of diversity for Hill numbers of different orders.

Then, we used generalized linear models (GLMs) to investigate the influence of elevation on the three facets of diversity for these Hill orders. Overall, for each diversity facet and Hill order, we fitted five models that included all possible combinations of effects: 1) no effect (*i.e.*, null model), 2) only elevation, 3) only transect, 4) similar effect of elevation across transects (*i.e.*, without interaction between elevation and transect or parallel slopes model), and 5) differential effect of elevation among transects (*i.e.*, with interaction between elevation and transect or different slopes model). We used a negative binomial error distribution (and a log-link function) to account for overdispersion. We compared alternative models based on the corrected Akaike information criterion (AICc). We considered the models within two AICc units of the model with the lowest AICc value. Among these models, we selected the most complex and calculated the explained deviance (*D*^2^) as a measure of goodness of fit for GLMs. Analyses were conducted using ‘MASS’ (Venables and Ripley 2002; v. 7.3-51.6) and ‘MuMIn’ (Barton 2018; v. 1.43.17) R packages.

All the R Statistical Software analyses were conducted using version 4.0.2 (R Core Team, 2020).

## 3 | RESULTS

Pearson′s correlations coefficients between number of trees per plot and any of the three facets of diversity (taxonomic, functional, and phylogenetic) for different Hill orders was very low, ranging between −0.22 and 0.08 and hence there was no need to consider the effect of sample size on diversity metrics.

When all transects were analysed together, taxonomic, and functional diversity were always significantly and positively correlated for *q* = 0, *q* = 1, and *q* = 2 (Fig. 2 and Appendix 1, Table A4). However, taxonomic, and phylogenetic diversity, and functional and phylogenetic diversity were significantly and positively correlated only for *q* = 0 and *q* = 1. After analysing the transects independently, we found the following.

- Taxonomic vs. functional diversity: always significantly and positively correlated for *q* = 0, *q* = 1, and *q* = 2.
- Taxonomic vs. phylogenetic diversity: significantly and positively correlated for *q* = 0 and *q* = 1 (except in Podocarpus for *q* = 1).
- Functional and phylogenetic diversity: significantly and positively correlated for *q* = 0 and *q* = 1 (except in Podocarpus for *q* = 1).

**FIGURE 2.**
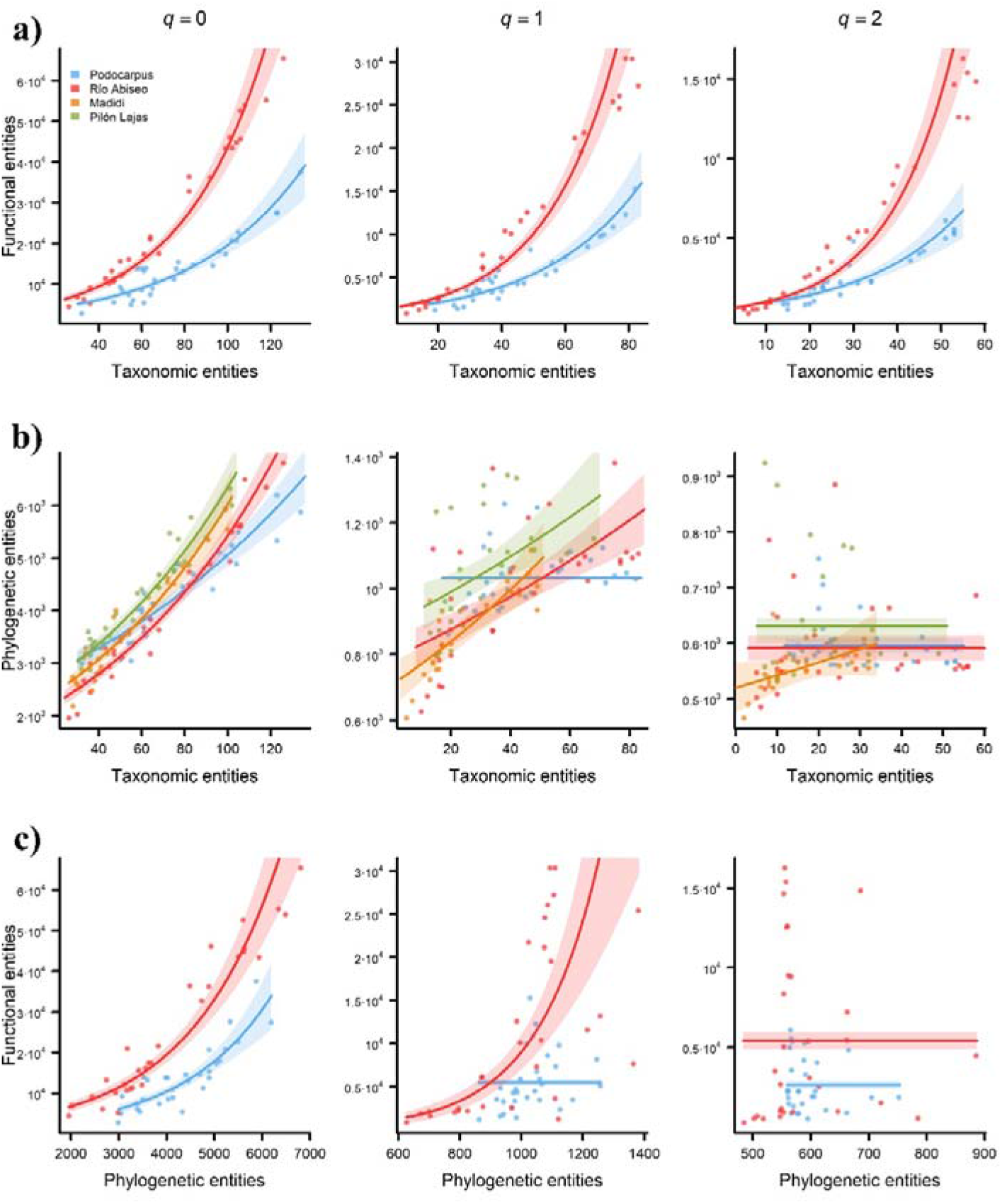
Pairwise comparisons of the three types of diversity (taxonomic, functional, and phylogenetic) for each of the four transects ranging elevational gradients in Andean tropical montane forests. Predictions of generalized linear models (GLMs) relating a) functional versus taxonomic diversity, b) phylogenetic versus taxonomic diversity, and c) functional versus phylogenetic diversity patterns for Hill orders of *q* = 0 (left-hand charts), *q* = 1 (central charts), and *q* = 2 (right-hand charts). Numbers represent the taxonomic, functional, and phylogenetic entities, a non-dimensional unit result of using the attribute diversity framework. Shaded area represents 95% confidence intervals based on the model predictions. GLMs were fitted using a negative binomial error distribution and a log-link function. Note that in certain charts we refrained from using the same scale in the axes to avoid them resulting visually uninformative.

Overall, results were consistent between transects. The three types of diversity values decreased from lower to higher elevations for *q* = 0, *q* = 1, and *q* = 2. This trend was strongest for *q* = 0 and weakest for *q* = 2, as indicated by both the slope of the regression curve (Fig. 3) and the explained deviance (Table 2). The only exception was phylogenetic diversity for *q* = 2, which appeared to increase slightly with the elevation. The best fitted models always included both elevation and transect (with or without interaction) as predictors, except for phylogenetic diversity for *q* = 2 (Fig. 3, Table 2 and Appendix 1, Table A5).

**FIGURE 3.**
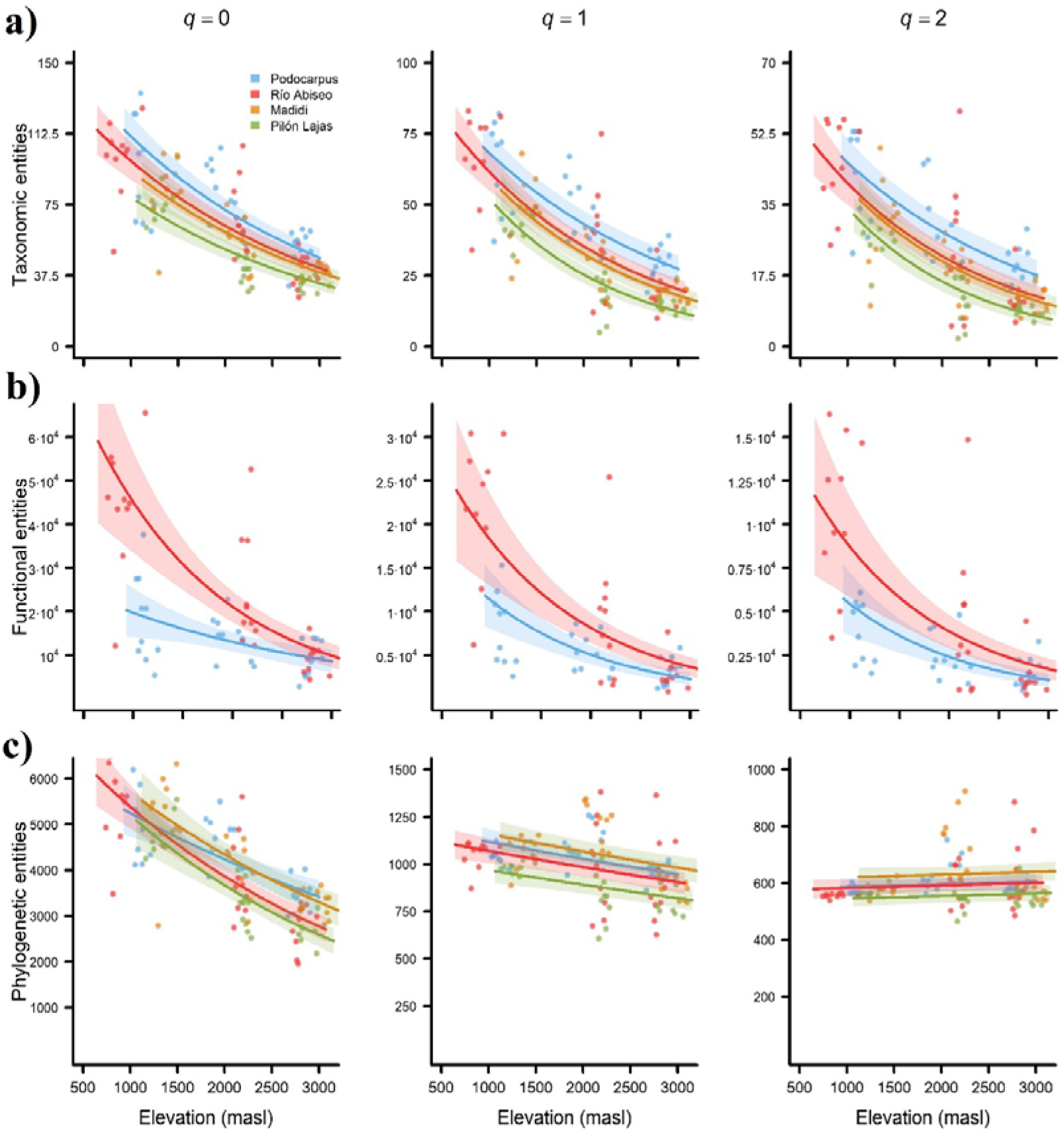
Patterns of taxonomic, functional, and phylogenetic diversity in Andean tropical montane forests along the four transects ranging elevational gradients. Predictions of generalized linear models (GLMs) relating a) taxonomic, b) functional, and c) phylogenetic diversity patterns with elevation (masl) for Hill orders of *q* = 0 (left-hand charts), *q* = 1 (central charts), and *q* = 2 (right-hand charts). Numbers represent the taxonomic, functional, and phylogenetic entities, a non-dimensional unit result of using the attribute diversity framework. Shaded area represents 95% confidence intervals based on the model predictions. GLMs were fitted using a negative binomial error distribution and a log-link function. Note that in certain charts we refrained from using the same scale in the axes to avoid them resulting visually uninformative.

## 4 | DISCUSSION

### 4.1 | Relationships between the three facets of diversity along elevation

The three facets of diversity clearly responded to elevation in a similar manner (Fig. 3), and they were significantly positive correlated in most cases (Fig. 2 and Appendix 1, Table A). These positive correlations may be expected due to changes in species richness along the elevational gradient, where more species are likely to exhibit more functional strategies and belong to more phylogenetic lineages (Losos 2008). However, this is not always necessarily true and empirical studies often show that two communities with equal number of species can differ greatly in terms of number of strategies and/or lineages (Devictor et al. 2010, Safi et al. 2011) and that a community exhibiting similar strategies may foster distantly related lineages (trait convergence) (Freckleton and Jetz 2009). Few studies have quantified the three types of diversity simultaneously along elevational gradients for plant communities (*e.g.*, Tanaka and Sato 2015, Chun and Lee 2017). In the tropical Andes, this type of study has only been conducted for vertebrates (*e.g.*, Dehling et al. 2014 for birds). Congruence between the three types of diversity along elevation has not been reported for plants, although pairwise correlations between two facets of diversity have been found: *e.g.*, functional and phylogenetic diversity (Tanaka and Sato 2015, Chun and Lee 2017). Regardless of the existence or absence of correlations between the different facets of diversity, all three provide complementary insights into the mechanisms that drive biodiversity patterns–as will be discussed in the following–, which could be obscured if only one is considered (Swenson 2013).

### 4.2 | Taxonomic diversity

Our results indicated a monotonic decrease in taxonomic diversity along elevation across all study transects that was consistent for different Hill orders (Fig. 3a), thereby suggesting that this decrease occurred regardless of whether taxa were weighted based on their abundance (*q* = 1 and *q* = 2) or not (*q* = 0).

General elevational decreases in taxonomic diversity of woody plants similar to those found in the present study have been widely reported (e.g., Kessler 2001, Nottingham et al. 2018), being the elevational decrease in temperature the ultimate cause of this trend (Grytnes and McCain 2007). However, studies have also detected a non-monotonic change in diversity along elevation, with a hump-shaped pattern where richness peaked at middle elevations (Gentry 1995, Rahbek 1995, Girardin et al. 2014). The existence of this hump in taxonomic diversity trend with elevation can be explained by tropical elevational gradients being characterized by a relatively stable condensation zone (cloud belt) at middle elevations, which provides favourable conditions for most organisms despite the lower temperatures relative to premontane elevations (Kessler and Kluge 2008, Huaraca Huasco et al. 2014). If this hump were to be present along our transects, our sampling design would have been unable to detect it because i) no sampling conducted below 1,000 masl (except for a very few plots in the Río Abiseo transect that did show an indication of a hump), and ii) a gap in our sampling around 1,5000 masl, (where the highest diversity could be expected) may have concealed a potential hump in species richness (Rahbek 2005, Nogués-Bravo et al. 2008).

### 4.3 | Functional diversity

Our results indicated a monotonic decrease in functional diversity along elevation across all transects and it was consistent for different Hill orders (Fig. 3b). Theory predicts that the range of successful functional strategies narrows along an environmental stress gradient (Cornwell and Ackerly 2009). In the case of elevation, the upslope increasingly harsher and more restrictive conditions mainly in terms of temperature and resource availability (Sundqvist et al. 2013) is expected to lead to the selective survival of taxa with conservative strategies –promoting storage and defence– that enable them to coping with the environment, and to the cull of these with acquisitive ones –enhancing photosynthesis and growth– (Poorter et al. 2009, Machac et al. 2011). This pattern has been detected previously in TMFs both for individual functional traits –including leaf size decreases (Salinas et al. 2011, Llerena-Zambrano et al. 2021) and wood density increases (Swenson and Enquist 2007, Slik et al. 2010)–, and for overall community functional diversity (Duivenvoorden and Cuello 2012, Schellenberger Costa et al. 2017). This evidence has been used in support of environmental filtering as an essential driver of plant community assembly (Swenson et al. 2012, Bañares-de-Dios et al. 2020).

The decrease in functional diversity was less pronounced for Hill numbers of higher order. Our results showed that Hill numbers that gave more weight to the most common entities (*q* = 2) accounted for two times less diversity than when the entities were strictly weighted by their abundance (*q* = 1) and four times less than when all entities received the same weight (*q* = 0) (Fig. 3b). These findings reflect how dominant/abundant species tend to be located at the core of the functional hyperspace whereas rare species tend to be peripheral. Dominant/abundant species exhibit strong traits-environment relationships enabling them a good performance in a given environment, thereby leading to greater demographic success. Conversely, rare species display sub-optimal traits –these species are more reliant on occasional migration events or stochastic environmental fluctuations–, which may explain their poorer demographic success (Umaña et al. 2017). In highly diverse ecosystems, such as tropical rainforests or coral reefs, rare species exhibit the less common traits (functional rarity), and thus disproportionately contribute to functional diversity, being irreplaceable and their loss not susceptible of being cushioned by the ecosystem high species richness as usually assumed. The reason is that the most common species have been shown to only support a relatively limited functionality (functional redundancy) (Mouillot et al. 2013, Leitão et al. 2016). Indeed, functional rarity sustains key ecosystem processes and services and provides an invaluable buffer against climate change and human disturbance (Cadotte et al. 2011, Violle et al. 2017). Therefore, the justification for conserving rare taxa should extend far beyond their taxonomic uniqueness, past, charisma or culture related arguments (Tucker et al. 2012).

### 4.4 | Phylogenetic diversity

Our results indicate that phylogenetic diversity decreased along elevation for *q* = 0 and *q* = 1 but increased slightly for *q* = 2 (Fig. 3c). In general, the overall decrease agrees with the predictions of the Tropical Niche Conservatism (TNC) hypothesis, which suggests that biological communities in the highlands phylogenetically are a subset of the communities found in the lowlands, and thus a decrease in phylogenetic diversity is expected with elevation (Wiens and Graham 2005). This pattern has been observed for other biota in Andean TMFs (*e.g.*, birds [Graham et al. 2009] or moths [Brehm et al. 2013]), but previous studies of Andean woody plants seem to give more support to the Out of the Tropical Lowlands (OTL) hypothesis (González-Caro et al. 2014, 2020, Tiede et al. 2016, Ramírez et al. 2019). Under OTL, the presence of temperate extratropical lineages within high elevation communities would be responsible for an increased phylogenetic alpha-diversity and turnover (Qian and Ricklefs 2016), which is the pattern we found when considering phylogenetic diversity for the most abundant species (*q* = 2).

Therefore, our results are not in full agreement with either hypothesis and endorse neither the support nor the rejection of the TNC or OTL (see also Tolmos et al. 2022). In this sense an alternative intermediate hypothesis termed Environmental Crossroads (Neves et al. 2020) has been proposed, which posits that floras from different origins – tropical and temperate– co-exist at middle elevations, at the edges –lower and upper, respectively– of their thermal tolerances (Griffiths et al. 2021). In any case, further research is needed, including determining the phylogenetic turnover and nestedness, as well as biogeographical origin of clades (Linan et al. 2020) and changes in their age along the elevational gradient (Qian 2014, Griffiths et al. 2020). A sampling schema that replicates transects along a broader latitudinal range, such as that used in the present study, would also help to assess the relative importance of the TNC vs. OTL. For example, several extratropical lineages such as *Quercus* prevail at high elevations in Colombia, which could have partly skewed the results reported by González-Caro et al. (2014, 2020) or Ramírez et al. (2019) based on studies that were latitudinally restricted to the northern Andes and which found evidence in agreement of OTL. *Quercus* is a Holarctic genus that migrated to South America in the last *ca.* 0.5 My (Hooghiemstra and Van Der Hammen 2004) and there may have not been sufficient time for it to reach the central Andes, where niche conservatism and the upslope migration of clades originated in tropical lowlands could have been the predominant mechanisms responsible for population by Andean flora.

In general, extratropical lineages that have migrated from temperate to tropical regions along the Andes, such as *Quercus* or *Alnus* in Colombia or *Araucaria* and *Nothofagus* in the southern Andes, are better adapted to cold conditions than strictly tropical lineages, and thus they are more likely to thrive in the uppermost TMF communities (Segovia et al. 2020). These taxa were absent from the transects that we studied but we found other floristic elements of Laurasian (*e.g.*, *Prunus* and *Cornus*) and Gondwanan (*e.g.*, *Podocarpus* and *Retrophyllum*) temperate origin, some of which tended to be dominant locally in the past and nowadays are far less abundant, with the only exception of certain fragments of relict forests (see Vicuña-Miñano 2005, Yaguana et al. 2012, Huamantupa-Chuquimaco et al. 2017). In this regard, the fact that our transects were located on isolated, protected areas of difficult access may have favoured the preservation of the native flora, allowing us to detect a slight increase in phylogenetic diversity at the highest elevations when giving more weigh to the most abundant taxa (*q* = 2) (Fig. 3c). Indeed, this slight increase resembles the phylogenetic pattern expected under the OTL. Therefore, as one moves far away from the central towards the northern or southern Andes, it would not be surprising to find an increase in the positive relationship between phylogenetic diversity and elevation, which may be detected by simply considering the relative abundances of taxa (*q* = 1) or equally considering all taxa (*q* = 0) given the greater prevalence of extratropical lineages.

Finally, it is important to note that the metrics used in this study to calculate phylogenetic diversity are strongly correlated with the number of lineages in recent evolutionary time but have weaker correlations with the number of lineages deeper in the evolutionary history of an assemblage (Dexter et al. 2019). Thus, our results might differ with the use of different metrics or the inclusion of different phylogenetic depths (Bose et al. 2019).

## 5 | CONCLUSIONS

We found an overall decrease in the three facets of diversity with elevation in Andean TMFs (the only exception was the phylogenetic diversity when we over-weighted dominant species). Our experimental design is particularly robust because we analysed four elevational gradients over a broad latitudinal range, thus avoiding the potential bias of other studies that focus on the much better known but geographically constrained northern Andean TMFs (Pérez-Escobar et al. 2022). Declines in taxonomic and functional diversity with elevation are consistent with the expected outcomes of an environmental filtering process in which temperature is the main driver of environmental variation. Differences in the steepness of the functional diversity trend between different Hill orders suggest that rare species play key and unique roles in the ecosystem, thereby supporting that the reasons for conserving rare taxa should extend far beyond their taxonomic or historical singularity. In addition, these findings suggest that niche conservatism and the upslope migration of tropical clades may be the predominant mechanisms responsible for shaping the flora of the central Andean highlands, given the limited arrival of temperate taxa in comparison to northern Andes. Further research across the entire Andean range, by integrating analyses of phylogenetic turnover and nestedness, as well as the biogeographical origin of clades and changes in their age along the elevational gradient will be important for elucidating the biogeographic history of the Andean slopes. The fact that the three types of diversity were positively correlated demonstrates that preserving TMF areas that harbour high taxonomic diversity will also preserve high functional and phylogenetic diversity in one of the most threatened biodiversity hotspots on Earth still poorly understood compared to tropical lowlands (ForestPlots.net 2021) and threatened by anthropogenic intervention and global warming (Salinas et al. 2021).

## ACKNOWLEDGEMENTS

G.B.D. was funded through grants from the Spanish Ministry of Education (FPU14/05303), Escuela Internacional de Doctorado - Universidad Rey Juan Carlos (Doctor Internacional 2017) and the Education and Research Department of Madrid Autonomous Region Government (REMEDINAL TE; S2018/EMT-4338). The study was supported through three grants from the Spanish Ministries of Economy and Competitiveness and Science and Technology (CGL2013-45634-P, CGL2016-75414-P, and PID2019-105064GB-I00), and a grant from Centro de Estudios de América Latina (CEAL) at Universidad Autónoma de Madrid and Banco Santander. We are indebted to those who helped with fieldwork: Stalin Japón, Wilson Remacha, Percy Malqui, “Rosho” Tamayo, Reinerio Ishuiza, Manuel Marca, Carlos Salas, José Sánchez, Anselmo Vergaray, Gonzalo Bañares, Ángel Delso, Julia González de Aledo, Mara Paneghel, Melecio Sullca, Raúl Huasurco, Honorato Pinto, Santiago Mamani, Aníbal Mamani, Armando Torrez, Justo Salas, Juan Mamani, Beto Apaza Coronel, Marcelo Reguerín, Óscar Quiroga, Juan Carlos Mamani, Rogelio Mamani, Vladimir Chura, Martín Chacón, and many others. We are especially thankful to Alex Nina, Jorge Armijos, Pablo Soliz, Leslie Cayola, Alfredo Fuentes, Peter Jørgensen and Tolentino Cueva, and the people of Los Alisos (Pataz, Peru) for their invaluable help. In addition, we are very grateful to Ministerio del Ambiente (MAE) in Ecuador, Servicio Nacional de Áreas Naturales Protegidas (SERNANP) in Peru, particularly Hugo Macedo, Vladimir Ramírez, Octavio Pecho, and Jhonny Ramos, and Ministerio de Medio Ambiente y Agua (MMAyA) in Bolivia. We extend our thanks to all the national park rangers who helped us, especially Rafael Galán, Guillermo Aguilar, Tito Heras, Percy Franco, Berardo Rojas, and Grover Benites. Permits to work in protected areas were granted by national authorities: Ecuador (MAE-DNB-CM-2015-0016; N° 001-2019-IC-FLO-FAU-DPAZCH-UPN-VS/MA), Peru (001-2016-SERNANP-PNRA-JEF), and Bolivia (MDRAyMA-VBRFMA-DGBAP-UAPVS N°2869/08).

## DATA AVAILABILITY STATEMENT

Plot census information for Ecuador (Podocarpus National Park) and Peru (Río Abiseo National Park) sites is available under request from the ForestPlots.net repository. Plot census information for Bolivia sites (Madidi National Park and Pilón-Lajas Biosphere Reserve) is available at the Tropicos database from Missouri Botanical Garden. Functional traits are stored in FunAndes (Báez et al., 2022), a database that integrates plant functional data from Andean forests, and they are in process of being incorporated to TRY database.

---

**Table A1.**
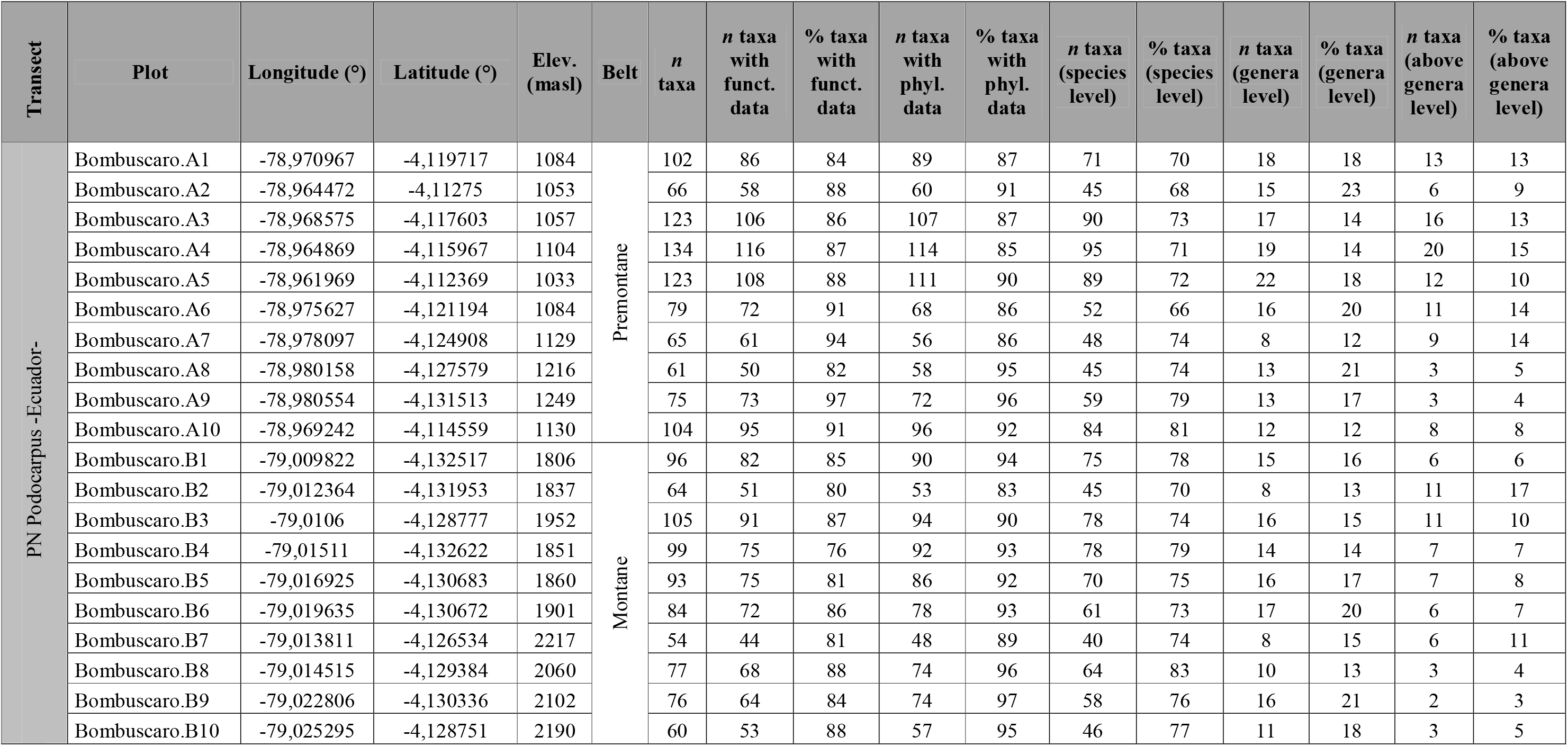
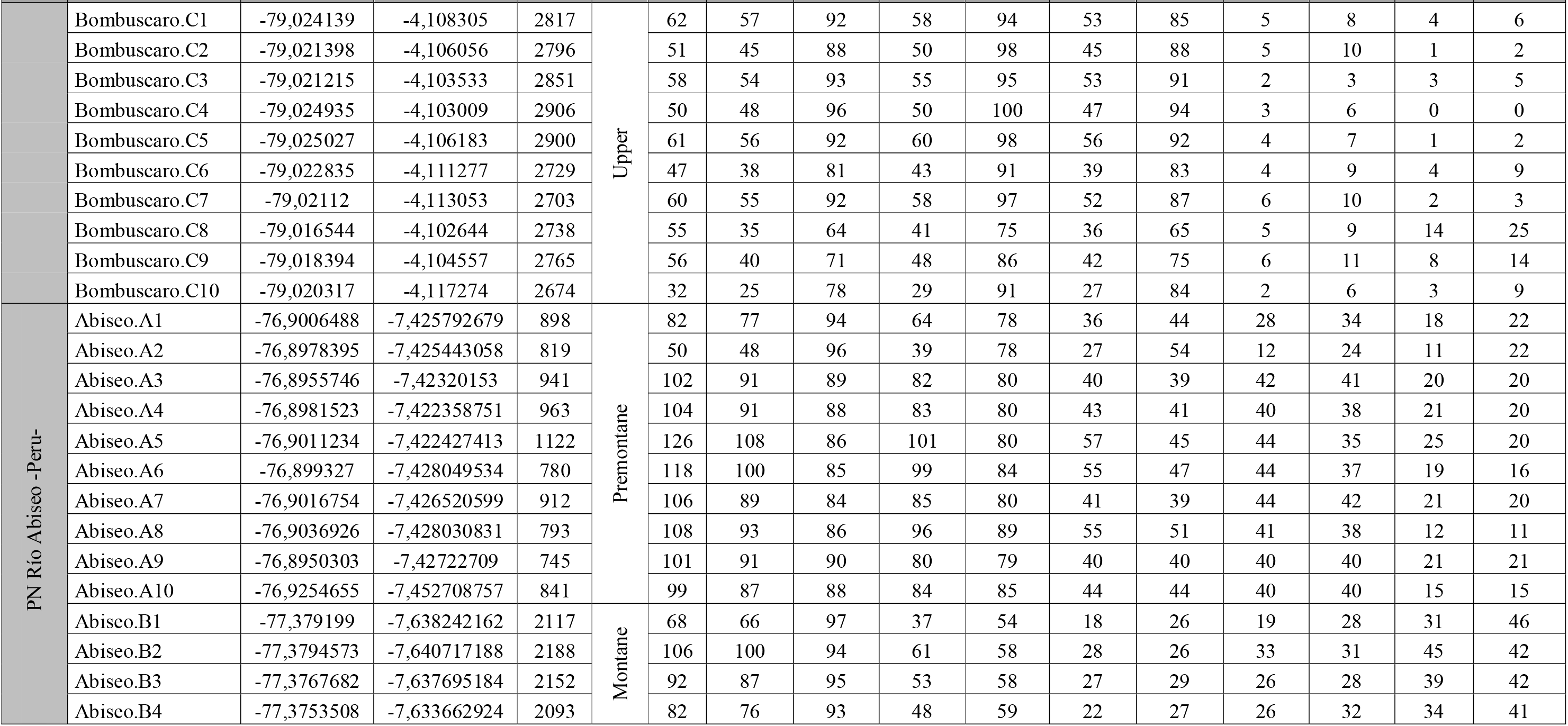
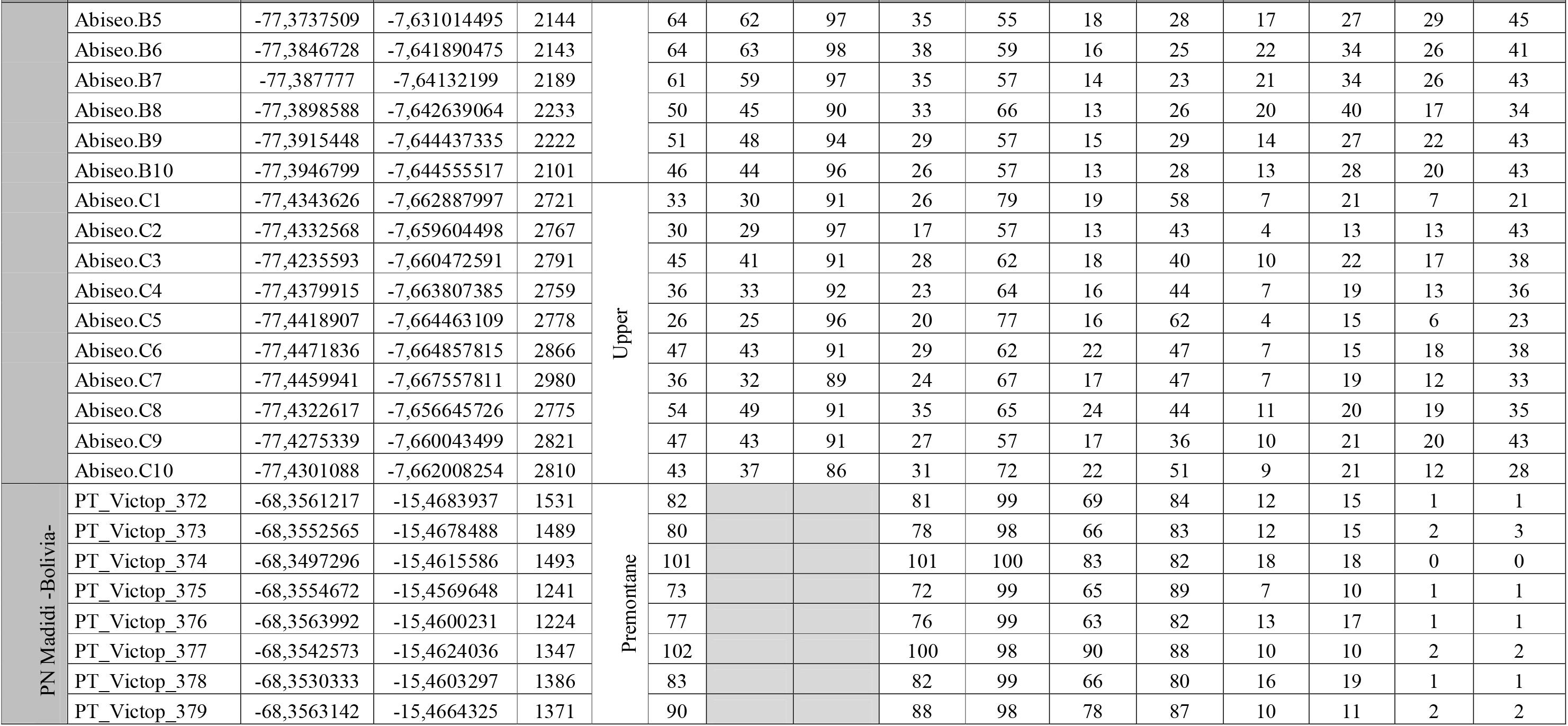
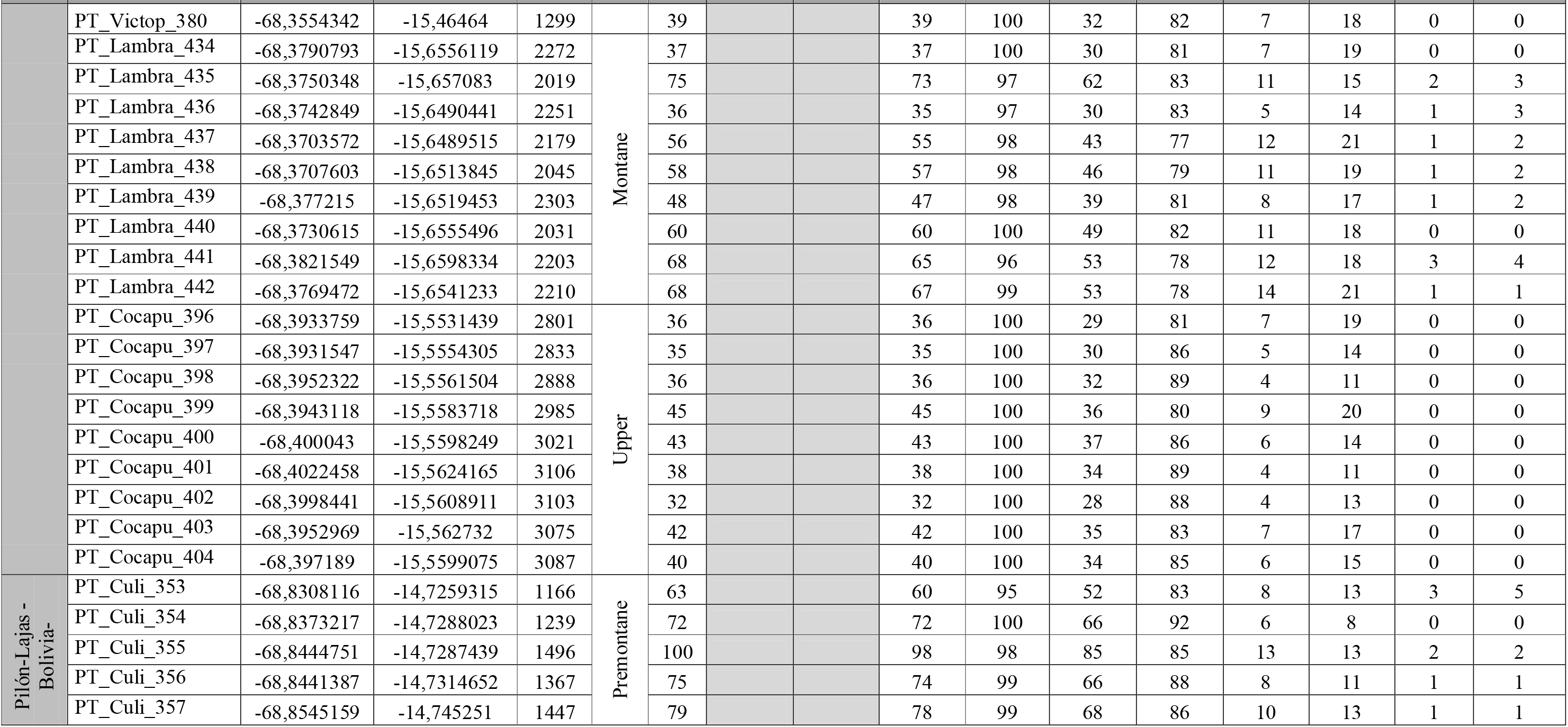
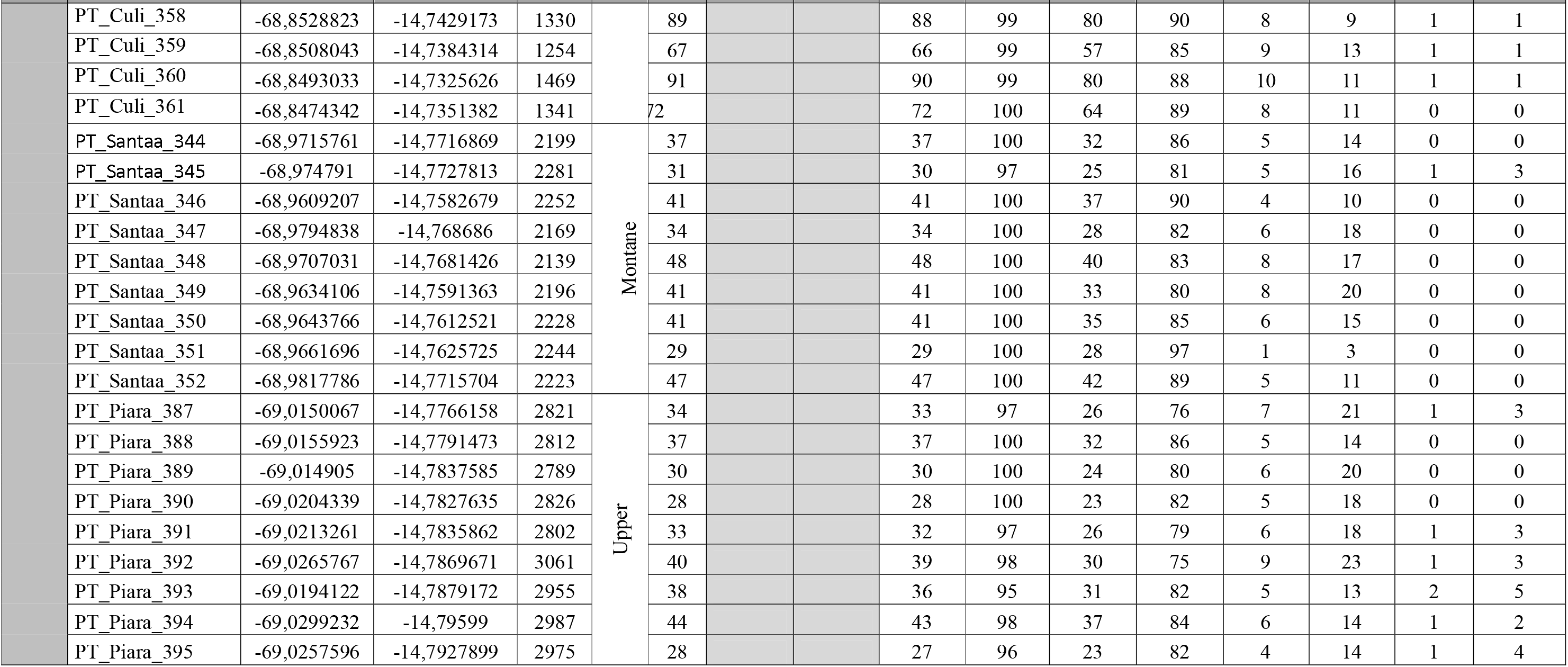
Information about the location, elevation, elevational belt, number of taxa, number and % of taxa over which functional and phylogenetic analysis were conducted, and taxonomic resolution of each plot of the four transects ranging elevational gradients. Light grey cells indicate lack of functional data for Madidi and Pilón-Lajas transects.

**Table A2.**
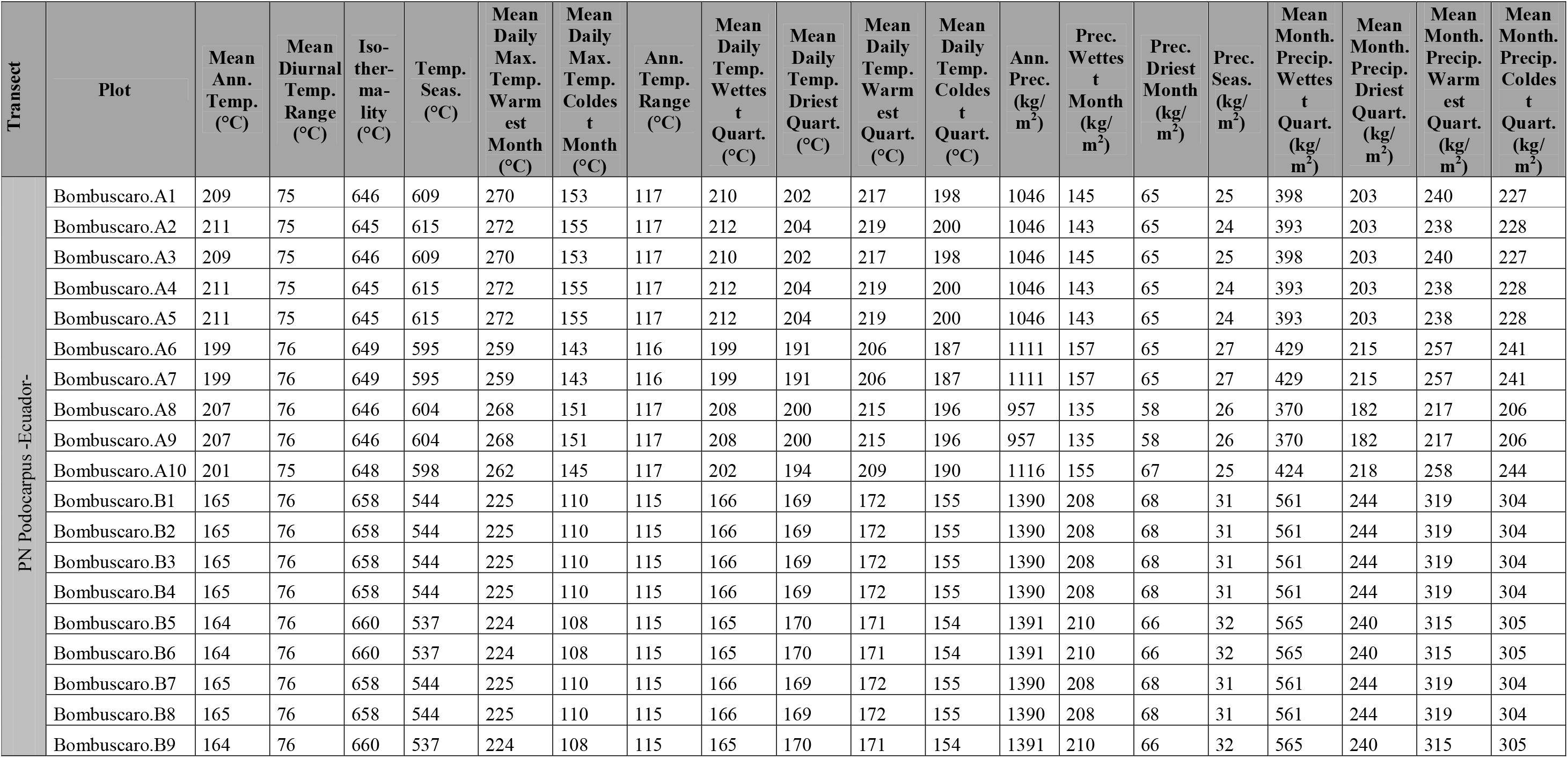
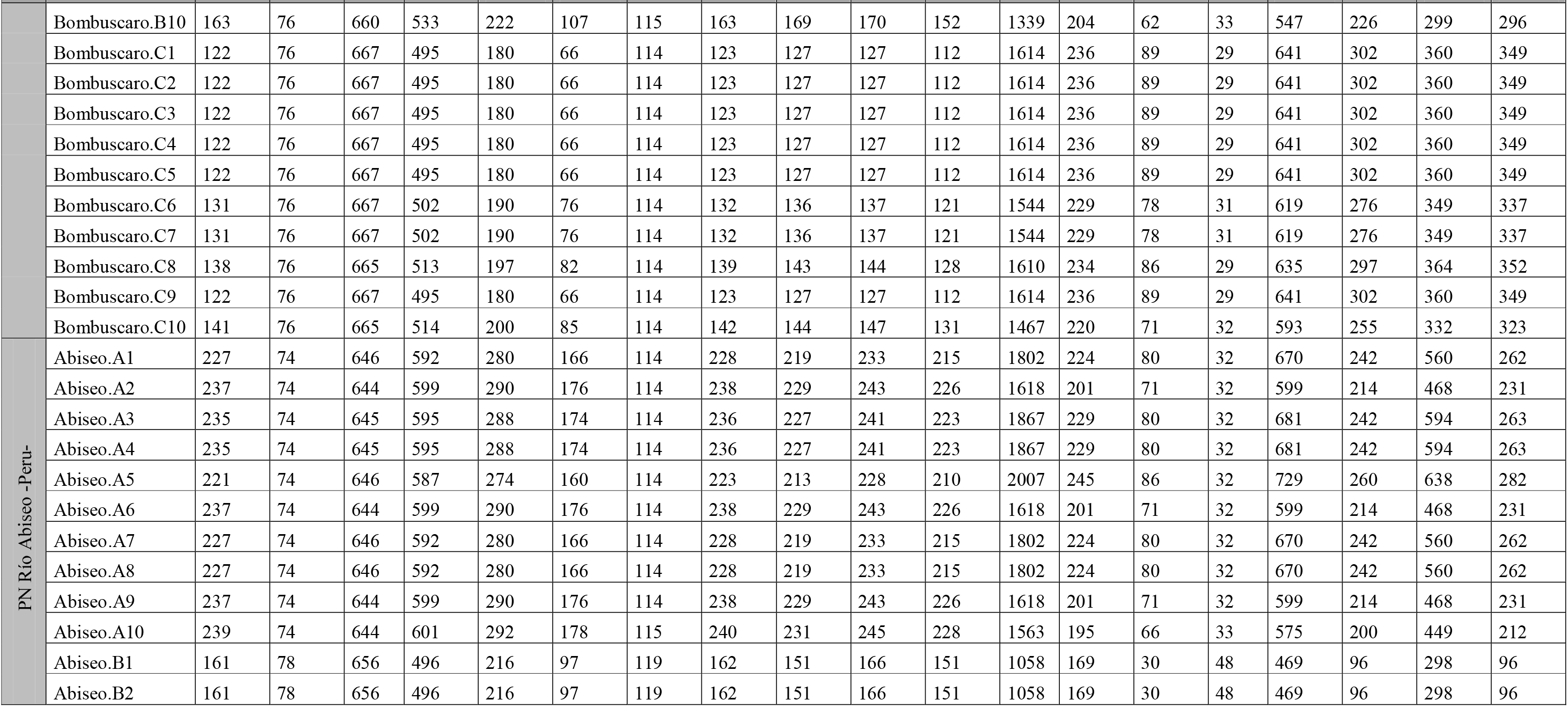
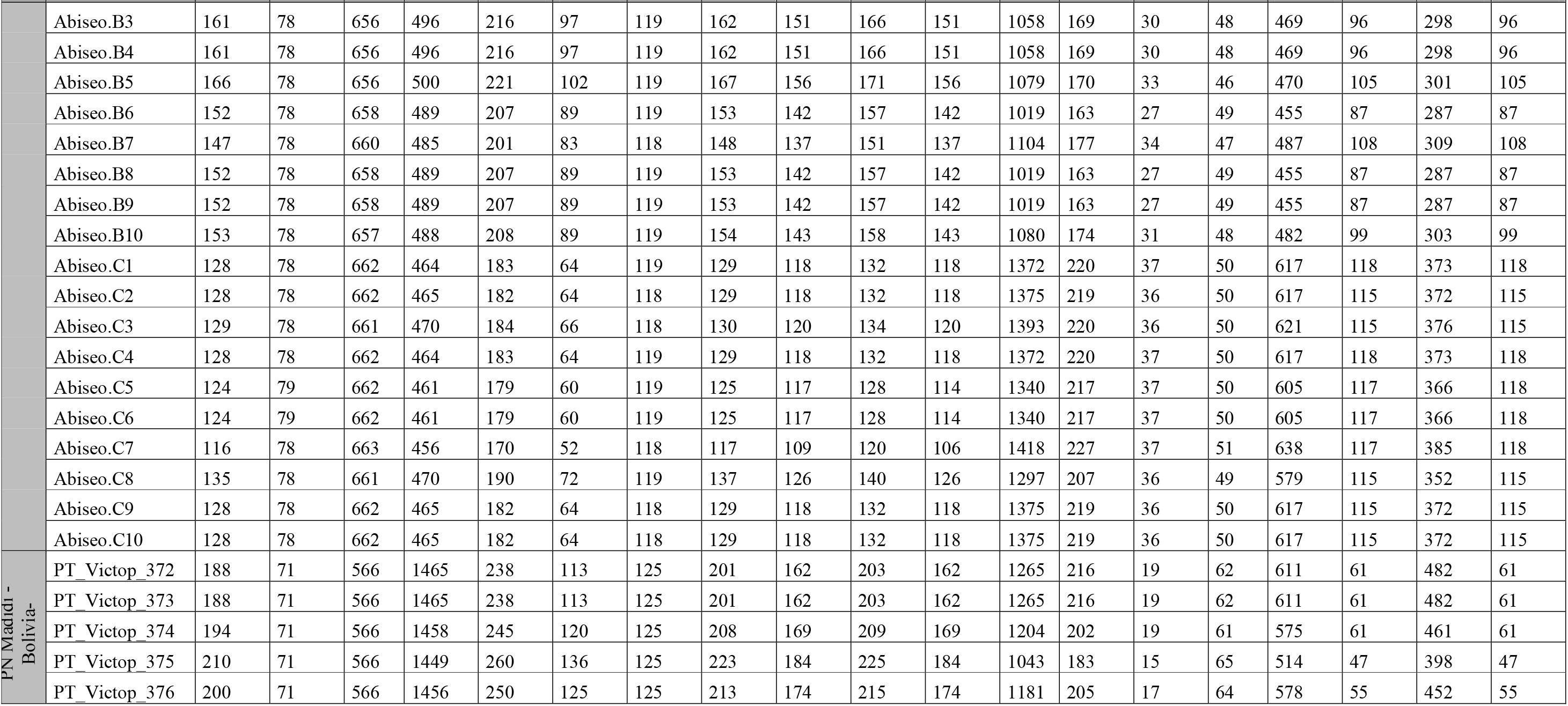
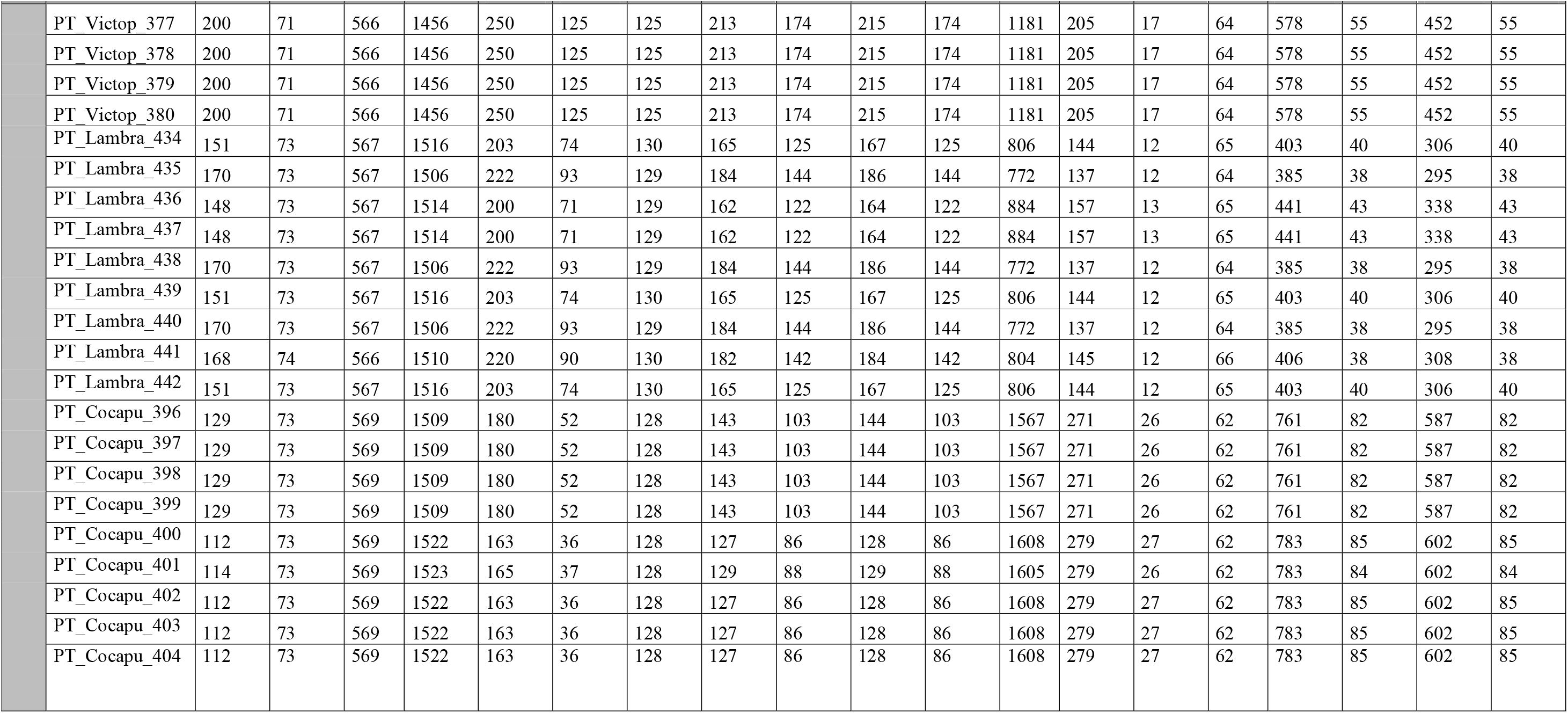
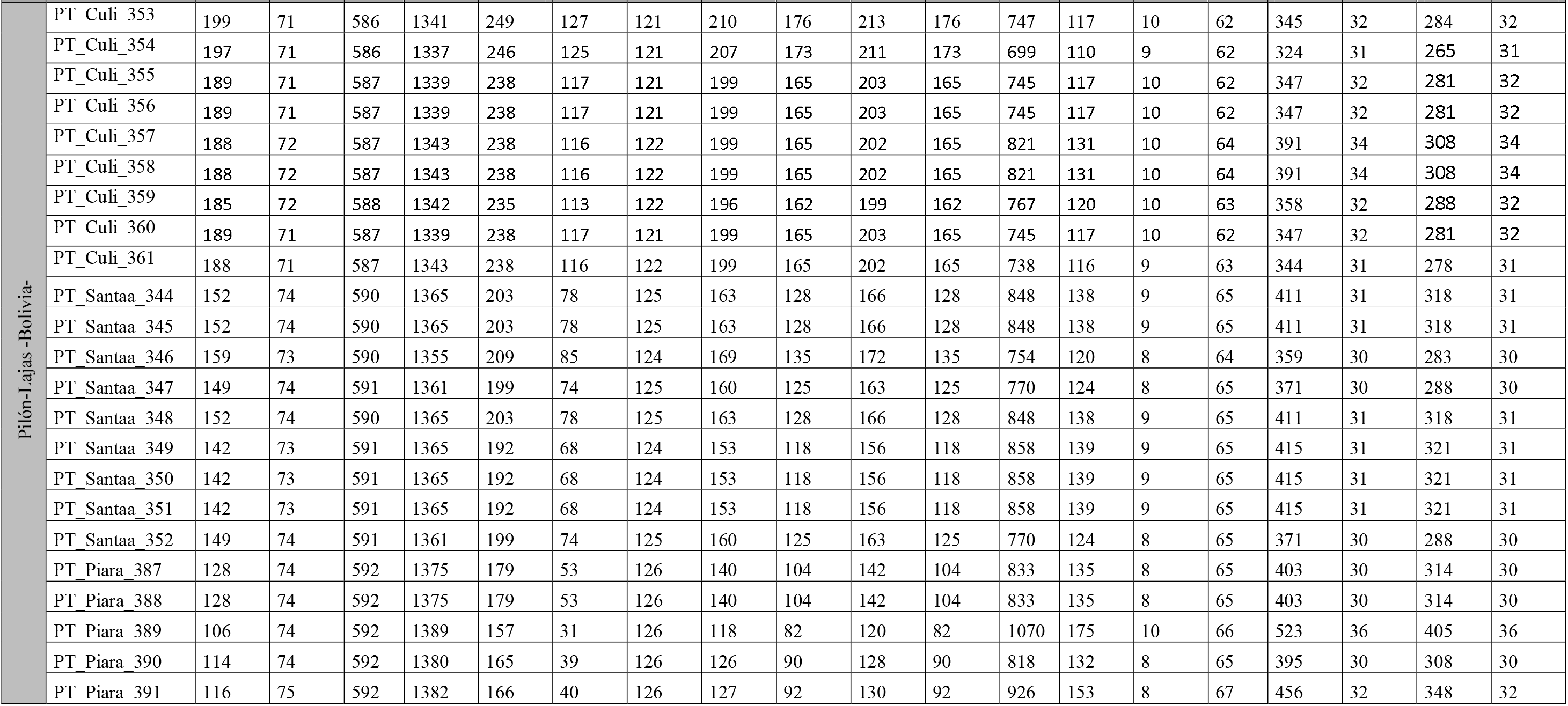
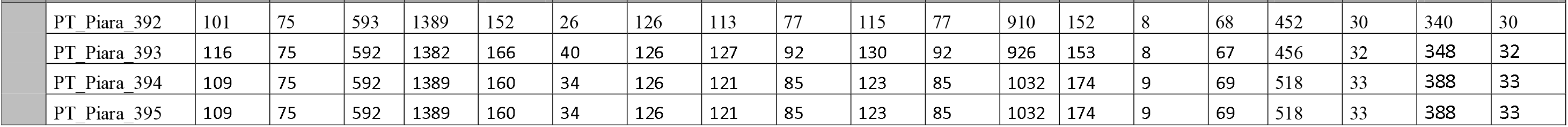
Bioclimatic variables values of each plot of the four transects ranging elevational gradients. Data obtained from CHELSA climatological dataset (Karger et al. 2017). Temp.: temperature, Seas.: seasonality, Quart.: quarter, Prec.: precipitation, Month.: monthly.

**Table A3.**
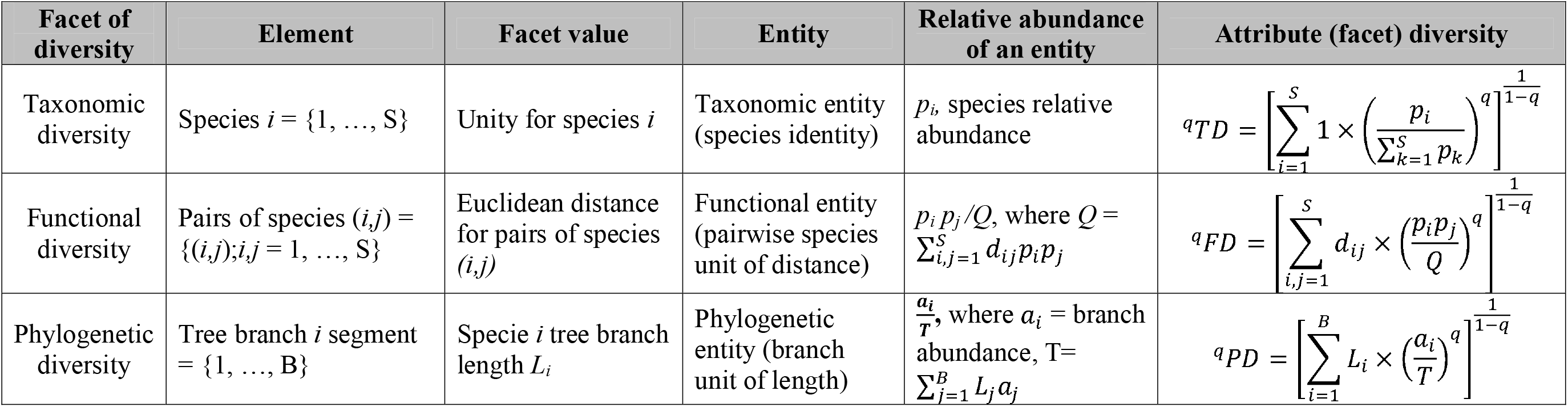
Unified framework for quantifying abundance-based diversity facets (taxonomic, functional, and phylogenetic) using generalized Hill numbers (with the parameter *q* defining the sensitivity to entities abundances). After Chao *et al*. (2014).

| Facet of diversity | Element | Facet value | Entity | Relative abundance of an entity | Attribute (facet) diversity |
| --- | --- | --- | --- | --- | --- |
| Taxonomic diversity | Species $i = \{1, \dots, S\}$ | Unity for species $i$ | Taxonomic entity (species identity) | $p_i$, species relative abundance | ${}^qTD = \left[ \sum_{i=1}^S 1 \times \left( \frac{p_i}{\sum_{k=1}^S p_k} \right)^q \right]^{\frac{1}{1-q}}$ |
| Functional diversity | Pairs of species $(i,j) = \{(i,j); i,j = 1, \dots, S\}$ | Euclidean distance for pairs of species $(i,j)$ | Functional entity (pairwise species unit of distance) | $p_i p_j / Q$, where $Q = \sum_{i,j=1}^S d_{ij} p_i p_j$ | ${}^qFD = \left[ \sum_{i,j=1}^S d_{ij} \times \left( \frac{p_i p_j}{Q} \right)^q \right]^{\frac{1}{1-q}}$ |
| Phylogenetic diversity | Tree branch $i$ segment = $\{1, \dots, B\}$ | Specie $i$ tree branch length $L_i$ | Phylogenetic entity (branch unit of length) | $\frac{a_i}{T}$, where $a_i$ = branch abundance, $T = \sum_{j=1}^B L_j a_j$ | ${}^qPD = \left[ \sum_{i=1}^B L_i \times \left( \frac{a_i}{T} \right)^q \right]^{\frac{1}{1-q}}$ |

**Table A4.**
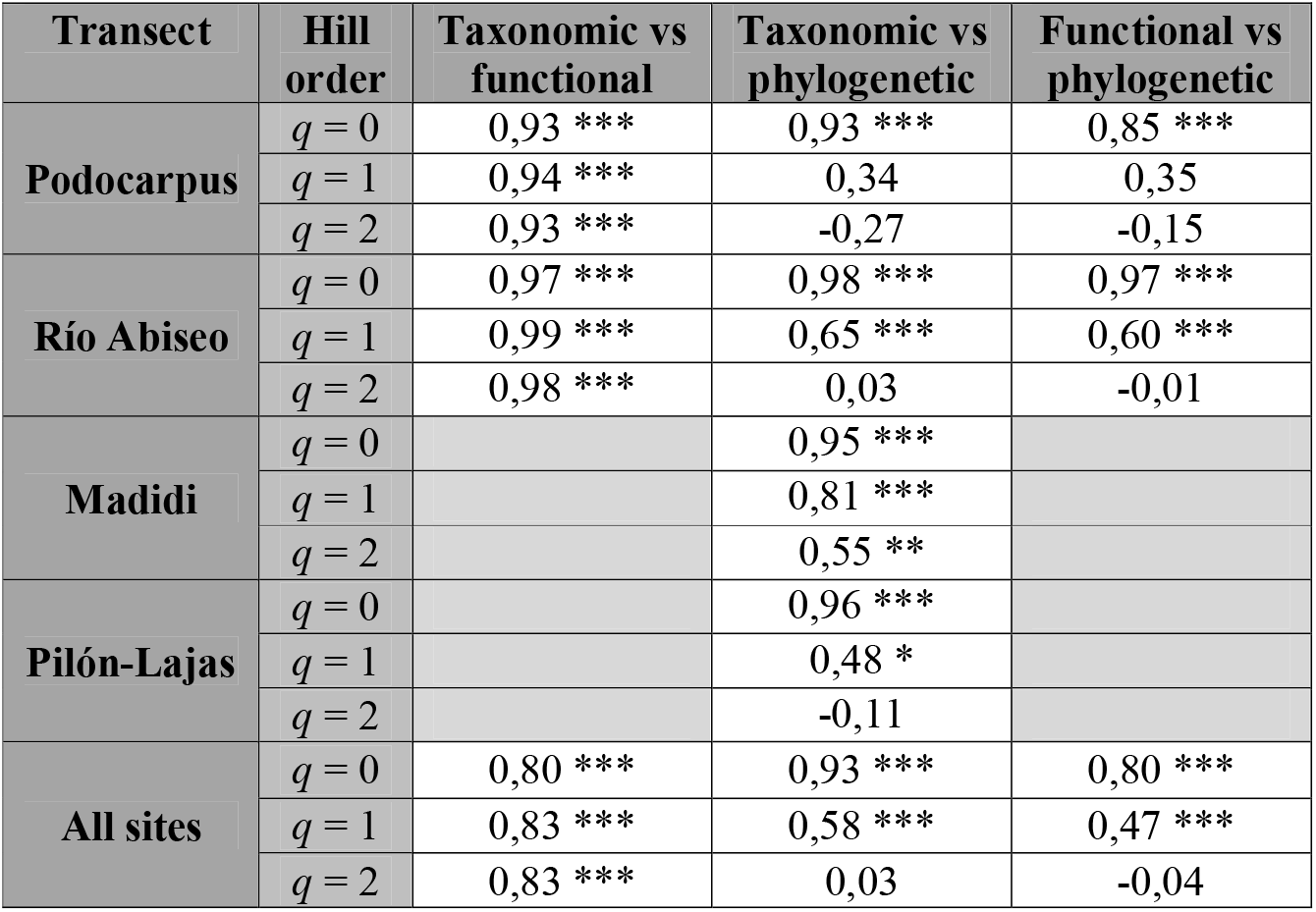
Pairwise Pearson’s correlations between the three facets of diversity (taxonomic, functional, and phylogenetic) for different Hill numbers’ orders. *: significant correlation (p<0.05); **: (p<0.01); ***: (p<0.001). Light grey cells indicate lack of functional data for Madidi and Pilón-Lajas transects.

**Table A5.**
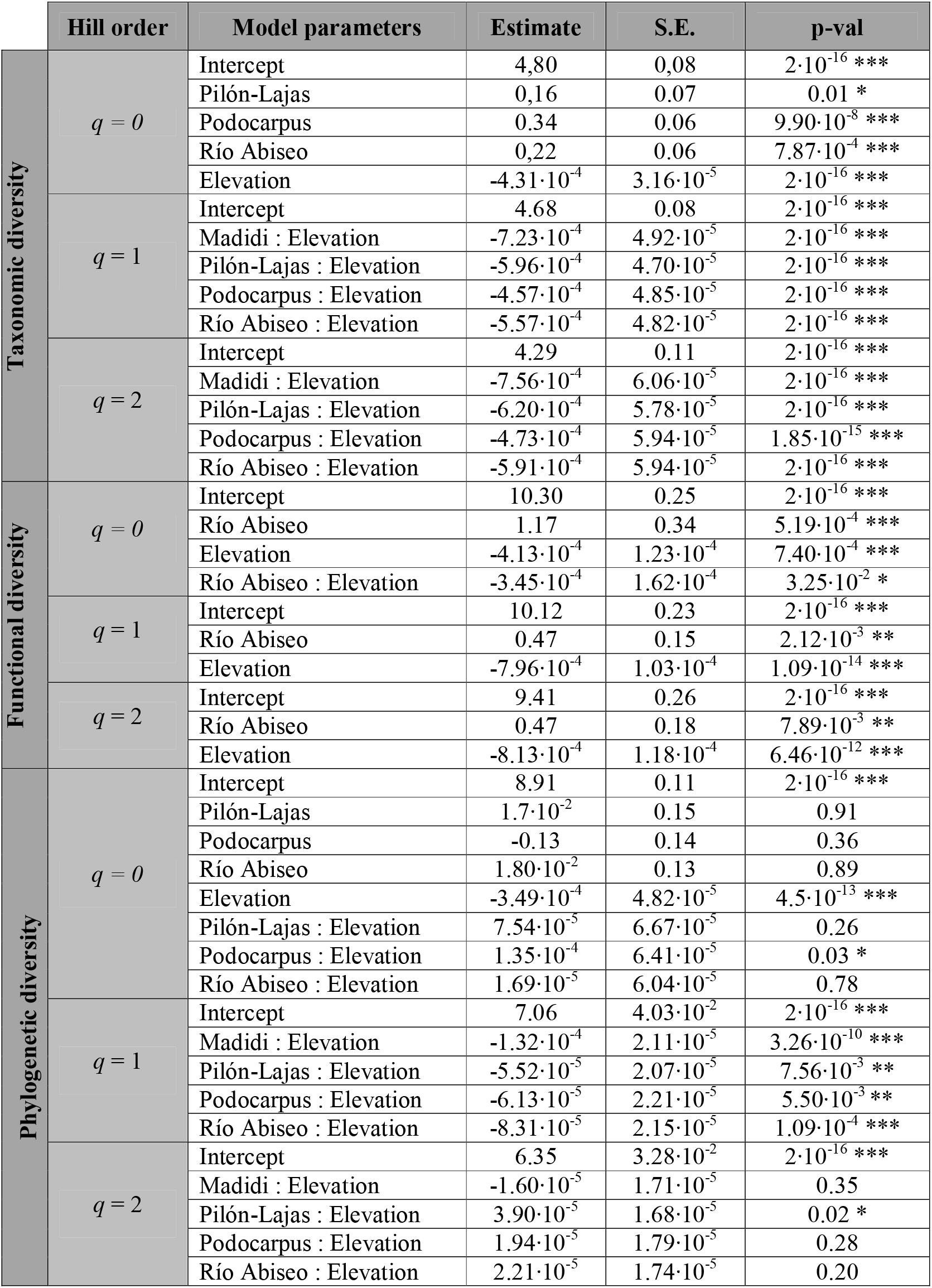
Model-averaged estimates, standard errors and p-values for selected variables in the best fitted models of taxonomic, functional, and phylogenetic diversity values calculated for Hill numbers of different order relative to distinct predictors for 114 plots of Andean TMFs. *: significant correlation (p<0.05), **: (p<0.01); ***: (p<0.001).

|  | Hill order | Model parameters | Estimate | S.E. | p-val |
| --- | --- | --- | --- | --- | --- |
| Taxonomic diversity | $q = 0$ | Intercept | 4,80 | 0,08 | $2 \cdot 10^{-16}$ *** |
|  |  | Pilón-Lajas | 0,16 | 0.07 | 0.01 * |
| | | Podocarpus | 0.34 | 0.06 | $9.90 \cdot 10^{-8}$ *** |
| | | Río Abiseo | 0,22 | 0.06 | $7.87 \cdot 10^{-4}$ *** |
| | | Elevation | $-4.31 \cdot 10^{-4}$ | $3.16 \cdot 10^{-5}$ | $2 \cdot 10^{-16}$ *** |
| | $q = 1$ | Intercept | 4.68 | 0.08 | $2 \cdot 10^{-16}$ *** |
| | | Madidi : Elevation | $-7.23 \cdot 10^{-4}$ | $4.92 \cdot 10^{-5}$ | $2 \cdot 10^{-16}$ *** |
| | | Pilón-Lajas : Elevation | $-5.96 \cdot 10^{-4}$ | $4.70 \cdot 10^{-5}$ | $2 \cdot 10^{-16}$ *** |
| | | Podocarpus : Elevation | $-4.57 \cdot 10^{-4}$ | $4.85 \cdot 10^{-5}$ | $2 \cdot 10^{-16}$ *** |
| | | Río Abiseo : Elevation | $-5.57 \cdot 10^{-4}$ | $4.82 \cdot 10^{-5}$ | $2 \cdot 10^{-16}$ *** |
| | $q = 2$ | Intercept | 4.29 | 0.11 | $2 \cdot 10^{-16}$ *** |
| | | Madidi : Elevation | $-7.56 \cdot 10^{-4}$ | $6.06 \cdot 10^{-5}$ | $2 \cdot 10^{-16}$ *** |
| | | Pilón-Lajas : Elevation | $-6.20 \cdot 10^{-4}$ | $5.78 \cdot 10^{-5}$ | $2 \cdot 10^{-16}$ *** |
| | | Podocarpus : Elevation | $-4.73 \cdot 10^{-4}$ | $5.94 \cdot 10^{-5}$ | $1.85 \cdot 10^{-15}$ *** |
| | | Río Abiseo : Elevation | $-5.91 \cdot 10^{-4}$ | $5.94 \cdot 10^{-5}$ | $2 \cdot 10^{-16}$ *** |
| Functional diversity | $q = 0$ | Intercept | 10.30 | 0.25 | $2 \cdot 10^{-16}$ *** |
| | | Río Abiseo | 1.17 | 0.34 | $5.19 \cdot 10^{-4}$ *** |
| | | Elevation | $-4.13 \cdot 10^{-4}$ | $1.23 \cdot 10^{-4}$ | $7.40 \cdot 10^{-4}$ *** |
| | | Río Abiseo : Elevation | $-3.45 \cdot 10^{-4}$ | $1.62 \cdot 10^{-4}$ | $3.25 \cdot 10^{-2}$ * |
| | $q = 1$ | Intercept | 10.12 | 0.23 | $2 \cdot 10^{-16}$ *** |
| | | Río Abiseo | 0.47 | 0.15 | $2.12 \cdot 10^{-3}$ ** |
| | | Elevation | $-7.96 \cdot 10^{-4}$ | $1.03 \cdot 10^{-4}$ | $1.09 \cdot 10^{-14}$ *** |
| | $q = 2$ | Intercept | 9.41 | 0.26 | $2 \cdot 10^{-16}$ *** |
| | | Río Abiseo | 0.47 | 0.18 | $7.89 \cdot 10^{-3}$ ** |
| | | Elevation | $-8.13 \cdot 10^{-4}$ | $1.18 \cdot 10^{-4}$ | $6.46 \cdot 10^{-12}$ *** |
| Phylogenetic diversity | $q = 0$ | Intercept | 8.91 | 0.11 | $2 \cdot 10^{-16}$ *** |
| | | Pilón-Lajas | $1.7 \cdot 10^{-2}$ | 0.15 | 0.91 |
|  |  | Podocarpus | -0.13 | 0.14 | 0.36 |
| | | Río Abiseo | $1.80 \cdot 10^{-2}$ | 0.13 | 0.89 |
| | | Elevation | $-3.49 \cdot 10^{-4}$ | $4.82 \cdot 10^{-5}$ | $4.5 \cdot 10^{-13}$ *** |
| | | Pilón-Lajas : Elevation | $7.54 \cdot 10^{-5}$ | $6.67 \cdot 10^{-5}$ | 0.26 |
| | | Podocarpus : Elevation | $1.35 \cdot 10^{-4}$ | $6.41 \cdot 10^{-5}$ | 0.03 * |
| | | Río Abiseo : Elevation | $1.69 \cdot 10^{-5}$ | $6.04 \cdot 10^{-5}$ | 0.78 |
| | $q = 1$ | Intercept | 7.06 | $4.03 \cdot 10^{-2}$ | $2 \cdot 10^{-16}$ *** |
| | | Madidi : Elevation | $-1.32 \cdot 10^{-4}$ | $2.11 \cdot 10^{-5}$ | $3.26 \cdot 10^{-10}$ *** |
| | | Pilón-Lajas : Elevation | $-5.52 \cdot 10^{-5}$ | $2.07 \cdot 10^{-5}$ | $7.56 \cdot 10^{-3}$ ** |
| | | Podocarpus : Elevation | $-6.13 \cdot 10^{-5}$ | $2.21 \cdot 10^{-5}$ | $5.50 \cdot 10^{-3}$ ** |
| | | Río Abiseo : Elevation | $-8.31 \cdot 10^{-5}$ | $2.15 \cdot 10^{-5}$ | $1.09 \cdot 10^{-4}$ *** |
| | $q = 2$ | Intercept | 6.35 | $3.28 \cdot 10^{-2}$ | $2 \cdot 10^{-16}$ *** |
| | | Madidi : Elevation | $-1.60 \cdot 10^{-5}$ | $1.71 \cdot 10^{-5}$ | 0.35 |
| | | Pilón-Lajas : Elevation | $3.90 \cdot 10^{-5}$ | $1.68 \cdot 10^{-5}$ | 0.02 * |
| | | Podocarpus : Elevation | $1.94 \cdot 10^{-5}$ | $1.79 \cdot 10^{-5}$ | 0.28 |
| | | Río Abiseo : Elevation | $2.21 \cdot 10^{-5}$ | $1.74 \cdot 10^{-5}$ | 0.20 |

## REFERENCES

Antonelli, A., W. D. Kissling, S. G. A. Flantua, M. A. Bermúdez, A. Mulch, A. N. Muellner-Riehl, H. Kreft, H. P. Linder, C. Badgley, J. Fjeldså, S. A. Fritz, C. Rahbek, F. Herman, H. Hooghiemstra, and C. Hoorn. 2018. Geological and climatic influences on mountain biodiversity. Nature Geoscience 11:718–725.

Antonelli, A., J. A. A. Nylander, C. Persson, and I. Sanmartín. 2009. Tracing the impact of the Andean uplift on Neotropical plant evolution 106:9749–9754.

Arellano, G., V. Cala, A. Fuentes, L. Cayola, P. M. Jorgensen, and M. J. Macía. 2016. A standard protocol for woody plant inventories and soil characterisation using temporary 0.1-Ha plots in tropical forests. Journal of Tropical Forest Science 28:508–516.

Barton, K. 2018. MuMIn: Multi-Model Inference. R package.

de Bello, F., S. Lavorel, S. Lavergne, C. H. Albert, I. Boulangeat, F. Mazel, and W. Thuiller. 2013. Hierarchical effects of environmental filters on the functional structure of plant communities: a case study in the French Alps. Ecography 36:393–402.

Bose R., B. R. Ramesh, R. Pélissier and F. Munoz. 2019. Phylogenetic diversity in the Western Ghats biodiversity hotspot reflects environmental filtering and past niche diversification of trees. Journal of Biogeography 46:145–157.

Brehm, G., P. Strutzenberger, and K. Fiedler. 2013. Phylogenetic diversity of geometrid moths decreases with elevation in the tropical Andes. Ecography 36:1247–1253.

Cadotte, M. W., K. Carscadden, and N. Mirotchnick. 2011. Beyond species: Functional diversity and the maintenance of ecological processes and services. Journal of Applied Ecology 48:1079–1087.

Caldas, F. 1803. Memoria sobre la nivelación de las plantas que se cultivan en la vecindad del Ecuador.

Cavender-Bares, J., K. H. Kozak, P. V. A. Fine, and S. W. Kembel. 2009. The merging of community ecology and phylogenetic biology. Ecology Letters 12:693–715.

Cayuela, L., I. Granzow-de la Cerda, F. S. Albuquerque, and D. J. Golicher. 2012. TAXONSTAND: An R package for species names standardisation in vegetation databases. Methods in Ecology and Evolution 3:1078–1083.

Chao, A., C.-H. Chiu, and L. Jost. 2010. Phylogenetic diversity measures based on Hill numbers. Philosophical Transactions of the Royal Society B: Biological Sciences 365:3599–3609.

Chao, A., C.-H. Chiu, and L. Jost. 2014a. Unifying Species Diversity, Phylogenetic Diversity, Functional Diversity, and Related Similarity and Differentiation Measures Through Hill Numbers. Annual Review of Ecology, Evolution, and Systematics 45:297–324.

Chao, A., N. J. Gotelli, T. Hsieh, E. L. Sander, K. Ma, R. K. Colwell, and A. M. Ellison. 2014b. Rarefaction and extrapolation with Hill numbers: a framework for sampling and estimation in species diversity studies. Ecological Monographs 84:45–67.

Chave, J., D. Coomes, S. Jansen, S. L. Lewis, N. G. Swenson, and A. E. Zanne. 2009. Towards a worldwide wood economics spectrum. Ecology Letters 12:351–366.

Chave, J., H. C. Muller-Landau, T. R. Baker, Ea, T. A. Easdale, H. Ter Steege, and C. O. Webb. 2006. Regional and phylogenetic variation of wood density across 2456 neotropical tree species. Ecological Applications 16:2356–2367.

Chiu, C. H., and A. Chao. 2014. Distance-based functional diversity measures and their decomposition: A framework based on Hill numbers. PLoS ONE 9:e100014.

Chun, J., and C. Lee. 2017. Disentangling the local-scale drivers of taxonomic, phylogenetic and functional diversity in woody plant assemblages along elevational gradients in South Korea. PLoS ONE 12:e0185763.

Cornelissen, J. H. C., S. Lavorel, E. Garnier, S. Díaz, N. Buchmann, D. E. Gurvich, P. B. Reich, H. Steege, H. D. Morgan, M. G. A. Van Der Heijden, J. G. Pausas, and H. Poorter. 2003. A handbook of protocols for standardised and easy measurement of plant functional traits worldwide. Australian Journal of Botany 51:335–380.

Cornwell, W. K., and D. D. Ackerly. 2009. Community assembly and shifts in the distribution of functional trait values across an environmental gradient in coastal California. Ecological Monographs 79:109–126.

Dani, R.S., P.K. Divakar, C.B. Baniya. 2023. Diversity and composition of plant species along elevational gradient: research trends. Biodiversity and Conservation 32: 2961–2980.

Dehling, D. M., S. A. Fritz, T. Töpfer, M. Päckert, P. Estler, K. Böhning-Gaese, and M. Schleuning. 2014. Functional and phylogenetic diversity and assemblage structure of frugivorous birds along an elevational gradient in the tropical Andes. Ecography 37:1047–1055.

Devictor, V., D. Mouillot, C. Meynard, F. Jiguet, W. Thuiller, and N. Mouquet. 2010. Spatial mismatch and congruence between taxonomic, phylogenetic and functional diversity: The need for integrative conservation strategies in a changing world. Ecology Letters 13:1030–1040.

Dexter, K. G., R A. Segovia and A. R. Griffiths. 2019. Exploring the Concept of Lineage Diversity across North American Forests. Forests 10(6):520.

Diaz, S., and M. Cabido. 1997. Plant functional types and ecosystem function in relation to global change. Journal of Vegetation Science 8:463–474.

Duivenvoorden, J. F., and N. L. Cuello. 2012. Functional trait state diversity of Andean forests in Venezuela changes with altitude. Journal of Vegetation Science 23:1105–1113.

Fadrique, B., and J. Homeier. 2016. Elevation and topography influence community structure, biomass and host tree interactions of lianas in tropical montane forests of southern Ecuador. Journal of Vegetation Science 27:958–968.

ForestPlots.net. 2021. Taking the pulse of Earth’s tropical forests using networks of highly distributed plots. Biological Conservation 108849.

Freckleton, R. P., and W. Jetz. 2009. Space versus phylogeny: Disentangling phylogenetic and spatial signals in comparative data. Proceedings of the Royal Society B: Biological Sciences 276:21–30.

Gentry, A. H. 1995. Patterns of diversity and floristic composition in Neotropical montane forests. Pages 103–126 in S. Churchill, H. Balsev, E. Forero, and J. Luteyn, editors. Biodiversity and conservation of Neotropical montane forests. The New York Botanical Garden, New York.

Girardin, C. A. J., W. Farfan-Rios, K. Garcia, K. J. Feeley, P. M. Jørgensen, A. A. Murakami, L. Cayola Pérez, R. Seidel, N. Paniagua, A. F. Fuentes Claros, C. Maldonado, M. Silman, N. Salinas, C. Reynel, D. A. Neill, M. Serrano, C. J. Caballero, M. de los A. La Torre Cuadros, M. J. Macía, T. J. Killeen, and Y. Malhi. 2014. Spatial patterns of above-ground structure, biomass and composition in a network of six Andean elevation transects. Plant Ecology and Diversity 7:161– 171.

González-Caro, S., Á. Duque, K. J. Feeley, E. Cabrera, J. Phillips, S. Ramirez, and A. Yepes. 2020. The legacy of biogeographic history on the composition and structure of Andean forests. Ecology 101:1–11.

González-Caro, S., M. N. Umaña, E. Álvarez, P. R. Stevenson, and N. G. Swenson. 2014. Phylogenetic alpha and beta diversity in tropical tree assemblages along regional-scale environmental gradients in northwest South America. Journal of Plant Ecology 7:145–153.

Graham, C. H., A. C. Carnaval, C. D. Cadena, K. R. Zamudio, T. E. Roberts, J. L. Parra, C. M. Mccain, R. C. K. Bowie, C. Moritz, S. B. Baines, C. J. Schneider, J. Vanderwal, C. Rahbek, K. H. Kozak, and N. J. Sanders. 2014. The origin and maintenance of montane diversity: Integrating evolutionary and ecological processes. Ecography 37:711–719.

Graham, C. H., J. Parra, C. Rahbek, and J. McGuire. 2009. Phylogenetic structure in tropical hummingbird communities. Proceedings of the National Academy of Sciences 106:19673–19678.

Griffiths, A. R., M. R. Silman, W. Farfan-Rios, K. J. Feeley, K. G. Cabrera, P. Meir, N. Salinas, R. A. Segovia, and K. G. Dexter. 2021. Evolutionary Diversity Peaks at Mid-Elevations Along an Amazon-to-Andes Elevation Gradient. Frontiers in Ecology and Evolution 9:1–10.

Griffiths, A. R., M. R. Silman, W. Farfán Rios, K. J. Feeley, K. García Cabrera, P. Meir, N. Salinas, and K. G. Dexter. 2020. Evolutionary heritage shapes tree distributions along an Amazon-to-Andes elevation gradient. Biotropica 53:38–50.

Grytnes, J.-A., and C. M. McCain. 2007. Elevational Trends in Biodiversity. Pages 1–8 in S. A. Levin, editor. Encyclopedia of Biodiversity. Elsevier.

Guo, Q., D. A. Kelt, Z. Sun, H. Liu, L. Hu, H. Ren, and J. Wen. 2013. Global variation in elevational diversity patterns. Scientific Reports 2013 3:1 3:1–7.

Hooghiemstra, H., and T. Van Der Hammen. 2004. Quaternary Ice-Age dynamics in the Colombian Andes: developing an understanding of our legacy. Philosophical Transactions of the Royal Society of London. Series B: Biological Sciences 359:173–181.

Huamantupa-Chuquimaco, I., M. Luza-Victorio, Miguel Alfaro-Curitumay, Lucero Ururi, W. Huaman-Arque, M. Pedraza, and M. Peralvo. 2017. Diversidad y Biomasa Arbórea en los Bosques Andinos del Santuario Nacional del Ampay, Apurímac – Perú. Q’Euña - Sociedad Botánica del Cusco 8:07–26.

Huaraca Huasco, W., C. A. J. Girardin, C. E. Doughty, D. B. Metcalfe, L. D. Baca, J. E. Silva-Espejo, D. G. Cabrera, L. E. O. C. Aragão, A. R. Davila, T. R. Marthews, L. P. Huaraca-Quispe, I. Alzamora-Taype, L. E. Mora, W. Farfán-Rios, K. G. Cabrera, K. Halladay, N. Salinas-Revilla, M. R. Silman, P. Meir, and Y. Malhi. 2014. Seasonal production, allocation and cycling of carbon in two mid-elevation tropical montane forest plots in the Peruvian Andes. Plant Ecology and Diversity 7:125–142.

Jablonski, D., R. Kaustuv, and J. W. Valentine. 2006. Out of the Tropics: Evolutionary Diversity Gradient. Science 314:102–106.

Janzen, D. H. 1967. Why mountain passes are higher in the tropics. The American Naturalist 101:233–249.

Jin, Y., and H. Qian. 2019. V.PhyloMaker: and R package that can generate very large phylogenies for vascular plants. Ecography 42:1353–1359.

Karger, D. N., O. Conrad, J. Böhner, T. Kawohl, H. Kreft, R. W. Soria-auza, N. E. Zimmermann, H. P. Linder, and M. Kessler. 2017. Climatologies at high resolution for the earth’ s land surface areas. Scientific Data 4:170122.

Kerkhoff, A. J., P. Moriarty, and M. Weiser. 2014. The latitudinal species richness gradient in the New World woody angiosperms is consistent with the tropical conservatism hypothesis. Proceedings of the National Academy of Sciences 111:8125–8130.

Kessler, M. 2001 Patterns of diversity and range size of selected plant groups along an elevational transect in the Bolivian Andes. Biodiversity and Conservation 10:1897–1921.

Kessler, M., and J. Kluge. 2008. Tropical mountain forest: Patterns and processes in a Biodiversity hotspot. Page (S. R. Gradstein, J. Homeier, and J. Kluge, Eds.). Biodiversity and Ecology Series 2, Univ. Göttingen, Göttingen.

Körner, C. 2004. Mountain biodiversity, its causes and function. Ambio 33:11–17.

Körner, C. 2007. The use of ‘altitude’ in ecological research. Trends in Ecology and Evolution 22:569–574.

de la Cruz-Amo, L., G. Bañares-de-Dios, V. Cala, Í. Granzow-de la Cerda, C. I. Espinosa, A. Ledo, N. Salinas, M. J. Macía, and L. Cayuela. 2020. Trade-Offs Among Aboveground, Belowground, and Soil Organic Carbon Stocks Along Altitudinal Gradients in Andean Tropical Montane Forests. Frontiers in Plant Science 11:106.

Latham, R., and R. E. Ricklefs. 1993. Global patterns of tree species in moist forests: energy-diversity theory does not account for variation in species richness. Oikos 67:325–333.

Leitão, R. P., J. Zuanon, S. Villéger, S. E. Williams, C. Baraloto, C. Fortunel, F. P. Mendonça, and D. Mouillot. 2016. Rare species contribute disproportionately to the functional structure of species assemblages. Proc R Soc B 283:20160084.

Li, D. 2018. hillR: Taxonomic, functional and phylogenetic diversity and similarity through Hill numbers. Journal of Open Source Software 3:1041.

Linan, A. G., J. A. Myers, C. E. Edwards, A. E. Zanne, S. A. Smith, G. Arellano, L. Cayola, W. Farfán-Ríos, A. F. Fuentes, K. García-Cabrera, S. González-Caro, M. I. Loza, M. J. Macía, Y. Malhi, B. Nieto-Ariza, N. Salinas, M. Silman, and J. S. Tello. 2020. The evolutionary assembly of forest communities along environmental gradients: recent diversification or sorting of pre-adapted clades? bioRxiv:424032.

Llerena-Zambrano, M., J. C. Ordoñez, L. D. Llambí, M. van der Sande, E. Pinto, L. Salazar, and F. Cuesta. 2021. Minimum temperature drives community leaf trait variation in secondary montane forests along a 3000-m elevation gradient in the tropical Andes. https://doi.org/10.1080/17550874.2021.1903604.

Lomolino, M. V. 2001. Elevation gradients of species-density: historical and prospective views. Global Ecology and Biogeography 10:3–13.

Losos, J. B. 2008. Phylogenetic niche conservatism, phylogenetic signal and the relationship between phylogenetic relatedness and ecological similarity among species. Ecology Letters 11:995–1003.

Machac, A., M. Janda, R. R. Dunn, and N. J. Sanders. 2011. Elevational gradients in phylogenetic structure of ant communities reveal the interplay of biotic and abiotic constraints on diversity. Ecography 34:364–371.

Malhi, Y., M. Silman, N. Salinas, M. Bush, P. Meir, and S. Saatchi. 2010. Introduction: Elevation gradients in the tropics: Laboratories for ecosystem ecology and global change research. Global Change Biology 16:3171–3175.

McCain, C. M. 2005. Elevational gradients in diversity of small mammals. Ecology 86:366–372.

McGill, B. J., B. J. Enquist, E. Weiher, and M. Westoby. 2006. Rebuilding community ecology from functional traits. Trends in Ecology and Evolution 21:178–185.

Mittermeier, R. A., N. Myers, C. G. Mittermeier, P. Gil, M. Hoffman, J. Pilgrim, and T. Brooks. 2005. Hotspots revisited: Earth’s biologically richest and most endangered terrestrial ecoregions. Conservation International, Washington DC.

Moles, A. T., D. D. Ackerly, C. O. Webb, J. C. Twiddle, J. B. Dickie, and M. Westoby. 2005. A brief history of seed size. Science 307:576–580.

Mouillot, D., D. R. Bellwood, C. Baraloto, J. Chave, R. Galzin, M. Harmelin-Vivien, M. Kulbicki, S. Lavergne, S. Lavorel, N. Mouquet, C. E. T. Paine, J. Renaud, and W. Thuiller. 2013. Rare Species Support Vulnerable Functions in High-Diversity Ecosystems. PLoS Biology 11:e1001569.

Myers, N., R. A. Mittermeier, C. G. Mittermeier, G. A. B. Fonseca, and J. Kent. 2000. Biodiversity hotspots for conservation priorities. Nature 403:853–858.

Neves, D. M., K. G. Dexter, T. R. Baker, F. Coelho de Souza, A. T. Oliveira-Filho, L. P. Queiroz, H. C. Lima, M. F. Simon, G. P. Lewis, R. A. Segovia, L. Arroyo, C. Reynel, J. L. Marcelo-Peña, I. Huamantupa-Chuquimaco, D. Villarroel, G. A. Parada, A. Daza, R. Linares-Palomino, L. V. Ferreira, R. P. Salomão, G. S. Siqueira, M. T. Nascimento, C. N. Fraga, and R. T. Pennington. 2020. Evolutionary diversity in tropical tree communities peaks at intermediate precipitation. Scientific Reports 10:1 10:1–7.

Nogués-Bravo, D., M. B. Araújo, T. Romdal, and C. Rahbek. 2008. Scale effects and human impact on the elevational species richness gradients. Nature 453:216–219.

Nottingham, A.T., N. Fierer, B.L. Turner, J. Whitaker, N.J. Ostle, N.P. McNamara, R.D. Bardgett, J.W. Leff, N. Salinas, M.R. Silman, L.E. Kruuk, P. Meir. 2018. Microbes follow Humboldt: temperature drives plant and soil microbial biodiversity patterns from the Amazon to the Andes. Ecology 99(11):2455–2466.

Oksanen, J., F. G. Blanchet, M. Friendly, R. Kindt, P. Legendre, D. McGlinn, P. R. Minchin, R. B. O’Hara, G. L. Simpson, P. Solymos, M. H. H. Stevens, E. Szoecs and H. Wagner. 2019. https://CRAN.R-project.org/package=vegan

Pavoine, S., and M. B. Bonsall. 2011. Measuring biodiversity to explain community assembly: A unified approach. Biological Reviews 86:792–812.

Pérez-Escobar, O. A., A. Zizka, M. A. Bermúdez, A. S. Meseguer, F. L. Condamine, C. Hoorn, H. Hooghiemstra, Y. Pu, D. Bogarín, L. M. Boschman, R. T. Pennington, A. Antonelli, and G. Chomicki. 2022. The Andes through time: evolution and distribution of Andean floras. Trends in Plant Science 27:364–378.

Poorter, H., Ü. Niinemets, L. Poorter, I. J. Wright, and R. Villar. 2009. Causes and consequences of variation in leaf mass per area (LMA): a metaLanalysis. New Phytologist 182:565–588.

Qian, H. 2014. Contrasting relationships between clade age and temperature along latitudinal versus elevational gradients for woody angiosperms in forests of South America. Journal of Vegetation Science 25:1208–1215.

Qian, H., and R. E. Ricklefs. 2016. Out of the Tropical Lowlands: Latitude versus Elevation. Trends in Ecology and Evolution 31:738–741.

R Core Team. (2021). R: A language and environment for statistical computing. R Foundation for Statistical Computing. https://www.r-project.org/

Rahbek, C. 1995. The elevational gradient of species richness: a uniform pattern? Ecography 18:200–205.

Rahbek, C. 2005. The role of spatial scale and the perception of large-scale species-richness patterns. Ecology Letters 8:224–239.

Rahbek, C., M. K. Borregaard, A. Antonelli, R. K. Colwell, B. G. Holt, D. Nogues-bravo, C. M. Ø. Rasmussen, K. Richardson, M. T. Rosing, R. J. Whittaker, and J. Fjeldså. 2019a. Building mountain biodiversity: Geological and evolutionary processes. Science 365:1114–1119.

Rahbek, C., M. K. Borregaard, R. K. Colwell, B. Dalsgaard, B. G. Holt, N. Morueta-Holme, D. Nogues-bravo, R. J. Whittaker, and J. Fjeldså. 2019b. Humboldt’s enigma: What causes global patterns of mountain biodiversity? Science 365:1108–1113.

Ramírez, S., S. González-Caro, J. Phillips, E. Cabrera, K. J. Feeley, and Á. Duque. 2019. The influence of historical dispersal on the phylogenetic structure of tree communities in the tropical Andes. Biotropica 51:500–508.

Ricklefs, R. E., and D. Schluter. 1993. Species diversity in ecological communities: historical and geographical perspectives. Page (R. E. Ricklefs and D. Schluter, Eds.). University of Chicago Press, Chicago.

Safi, K., M. V. Cianciaruso, R. D. Loyola, D. Brito, K. Armour-Marshall, and J. A. F. Diniz-Filho. 2011. Understanding global patterns of mammalian functional and phylogenetic diversity. Philosophical Transactions of the Royal Society B: Biological Sciences 366:2536–2544.

Salinas, N., E. G. Cosio, M. Silman, P. Meir, A. T. Nottingham, R. M. Roman-Cuesta, and Y. Malhi. 2021. Editorial: Tropical Montane Forests in a Changing Environment. Frontiers in Plant Science 12:1652.

Salinas, N., Y. Malhi, P. Meir, M. Silman, R. Roman Cuesta, J. Huaman, D. Salinas, V. Huaman, A. Gibaja, M. Mamani, and F. Farfan. 2011. The sensitivity of tropical leaf litter decomposition to temperature: Results from a large-scale leaf translocation experiment along an elevation gradient in Peruvian forests. New Phytologist 189:967–977.

Sanders, N. J., and C. Rahbek. 2012. The patterns and causes of elevational diversity gradients. Ecography 35:1–3.

Sanmartín, I. 2012. Historical Biogeography: Evolution in Time and Space. Evo Edu Outreach 5, 555–568.

Schellenberger Costa, D., F. Gerschlauer, H. Pabst, A. Kühnel, B. Huwe, R. Kiese, Y. Kuzyakov, and M. Kleyer. 2017. Community-weighted means and functional dispersion of plant functional traits along environmental gradients on Mount Kilimanjaro. Journal of Vegetation Science 28:684–695.

Segovia, R. A., R. T. Pennington, T. R. Baker, F. C. de Souza, D. M. Neves, C. C. Davis, J. J. Armesto, A. T. Olivera-Filho, and K. G. Dexter. 2020. Freezing and water availability structure the evolutionary diversity of trees across the Americas. Science Advances 6.

Slik, J. W. F., S. I. Aiba, F. Q. Brearley, C. H. Cannon, O. Forshed, K. Kitayama, H. Nagamasu, R. Nilus, J. Payne, G. Paoli, A. D. Poulsen, N. Raes, D. Sheil, K. Sidiyasa, E. Suzuki, and J. L. C. H. van Valkenburg. 2010. Environmental correlates of tree biomass, basal area, wood specific gravity and stem density gradients in Borneo’s tropical forests. Global Ecology and Biogeography 19:50–60.

Sundqvist, M. K., N. J. Sanders, and D. A. Wardle. 2013. Community and Ecosystem Responses to Elevational Gradients: Processes, Mechanisms, and Insights for Global Change. Annual Review of Ecology, Evolution, and Systematics 44:261–280.

Swenson, N. G. 2013. The assembly of tropical tree communities - the advances and shortcomings of phylogenetic and functional trait analyses. Ecography 36:264– 276.

Swenson, N. G., and B. J. Enquist. 2007. Ecological and evolutionary determinants of a key plant functional trait, wood density and its community wide variation across latitude and elevation. American Journal of Botany 94:451–459.

Swenson, N. G., B. J. Enquist, J. Pither, A. J. Kerkhoff, B. Boyle, M. D. Weiser, J. J. Elser, W. F. Fagan, J. Forero-Montaña, N. Fyllas, N. J. B. Kraft, J. K. Lake, A. T. Moles, S. Patiño, O. L. Phillips, C. A. Price, P. B. Reich, C. A. Quesada, J. C. Stegen, R. Valencia, I. J. Wright, S. J. Wright, S. Andelman, P. M. Jørgensen, T. E. Lacher, A. Monteagudo, M. P. Núñez-Vargas, R. Vasquez-Martínez, and K. M. Nolting. 2012. The biogeography and filtering of woody plant functional diversity in North and South America. Global Ecology and Biogeography 21:798–808.

Tanaka, T., and T. Sato. 2015. Taxonomic, phylogenetic and functional diversities of ferns and lycophytes along an elevational gradient depend on taxonomic scales. Plant Ecology 216:1597–1609.

Thiers, B. (n.d.). Index Herbariorum. http://sweetgum.nybg.org/science/ih/.

Tiede, Y., J. Homeier, N. Cumbicus, J. Peña, J. Albrecht, B. Ziegenhagen, J. Bendix, R. Brandl, and N. Farwig. 2016. Phylogenetic niche conservatism does not explain elevational patterns of species richness, phylodiversity and family age of tree assemblages in andean rainforest. Erdkunde 70:83–106.

Tito, R., H. L. Vasconcelos, and K. J. Feeley. 2020. Mountain Ecosystems as Natural Laboratories for Climate Change Experiments. Frontiers in Forests and Global Change 3:1–8.

Tolmos, M. L., H. Kreft, J. Ramirez, R. Ospina, and D. Craven. 2022. Water and energy availability mediate biodiversity patterns along an elevational gradient in the tropical Andes. Journal of Biogeography 49:712–726.

Tucker, C. M., M. W. Cadotte, T. J. Davies, and T. G. Rebelo. 2012. Incorporating Geographical and Evolutionary Rarity into Conservation Prioritization. Conservation Biology 26:593–601.

Umaña, M. N., C. Zhang, M. Cao, L. Lin, and N. G. Swenson. 2017. A core-transient framework for trait-based community ecologyL: an example from a tropical tree seedling community. Ecology Letters 20:619–628.

Venables, W., and B. Ripley. 2002. Modern applied statistics with S. Springer, New York.

Vicuña-Miñano, E. E. 2005. Las Podocarpáceas de los bosques montanos del noroccidente peruano. Revista Peruana de Biologia 12:283–288.

Violle, C., W. Thuiller, N. Mouquet, F. Munoz, N. J. B. Kraft, M. W. Cadotte, S. W. Livingstone, and D. Mouillot. 2017. Functional Rarity: The Ecology of Outliers. Trends in Ecology and Evolution 32:356–367.

Wiens, J., and M. J. Donoghue. 2004. Historical biogeography, ecology and species richness. Trends in Ecology and Evolution 19:639–644.

Wiens, J., and C. H. Graham. 2005. Niche conservatism: integrating evollution, ecology and conservation biology. Annual Review of Ecology, Evolution, and Systematics 36:519–539.

Wright, I. J., P. B. Reich, M. Westoby, D. D. Ackerly, Z. Baruch, F. Bongers, J. Cavender-Bares, T. Chapin, J. H. C. Cornellssen, M. Diemer, J. Flexas, E. Garnier, P. K. Groom, J. Gulias, K. Hikosaka, B. B. Lamont, T. Lee, W. Lee, C. Lusk, J. J. Midgley, M. L. Navas, Ü. Niinemets, J. Oleksyn, H. Osada, H. Poorter, P. Pool, L. Prior, V. I. Pyankov, C. Roumet, S. C. Thomas, M. G. Tjoelker, E. J. Veneklaas, and R. Villar. 2004. The worldwide leaf economics spectrum. Nature 428:821–827.

Yaguana, C., D. Lozano, D. Neill, and M. Asanza. 2012. Diversidad florística y estructura del bosque nublado del Río Numbala, Zamora-Chinchipe, Ecuador. Revista Amazónica Ciencia y Tecnología 1:226–247.

