## Supplementary figures and images for "Woody plant taxonomic, functional, and phylogenetic diversity decrease along elevational gradients in Andean tropical montane forests: environmental filtering and arrival of temperate taxa"

### Appendix 1

Figure 1. Phylogenetic tree.

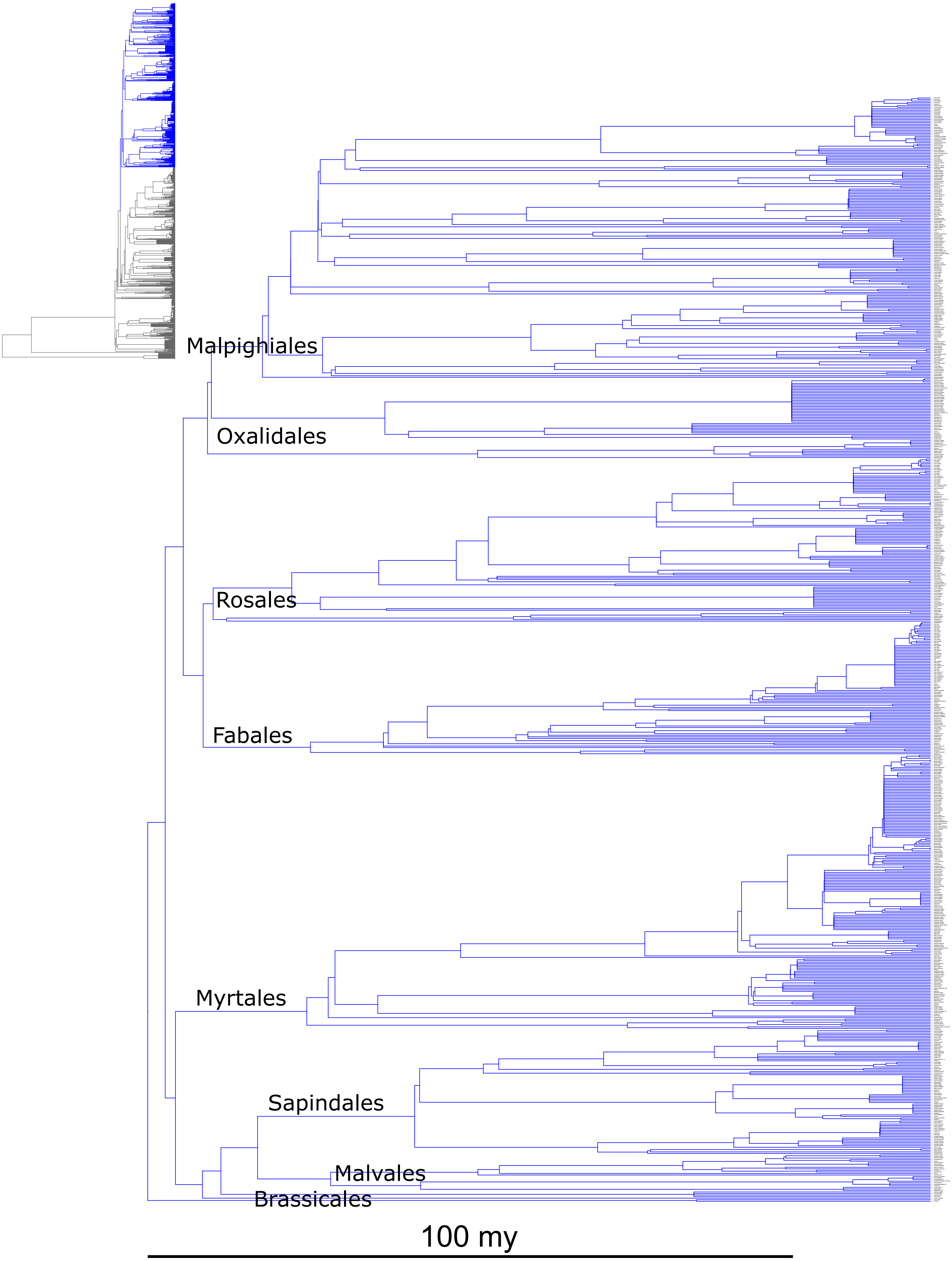

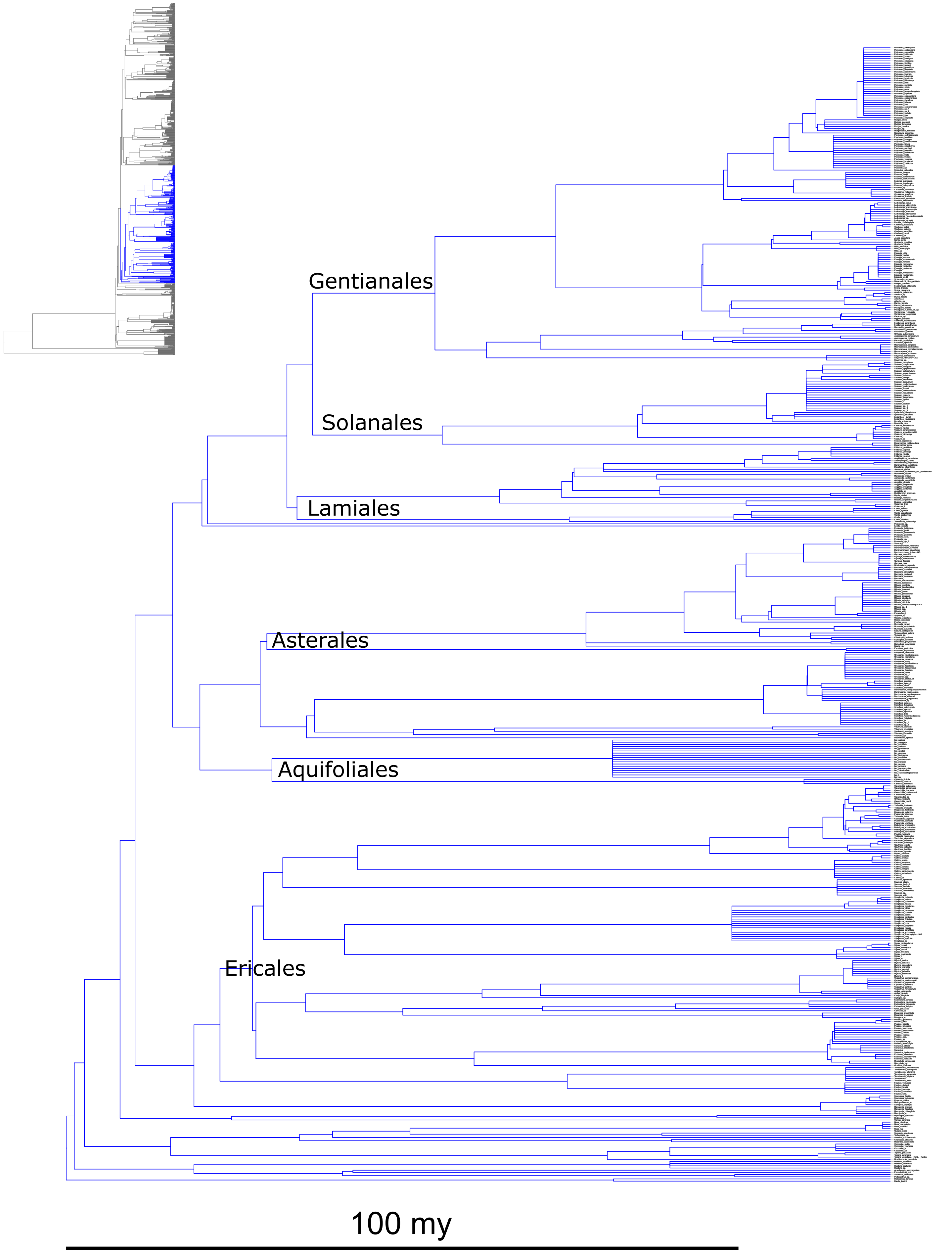

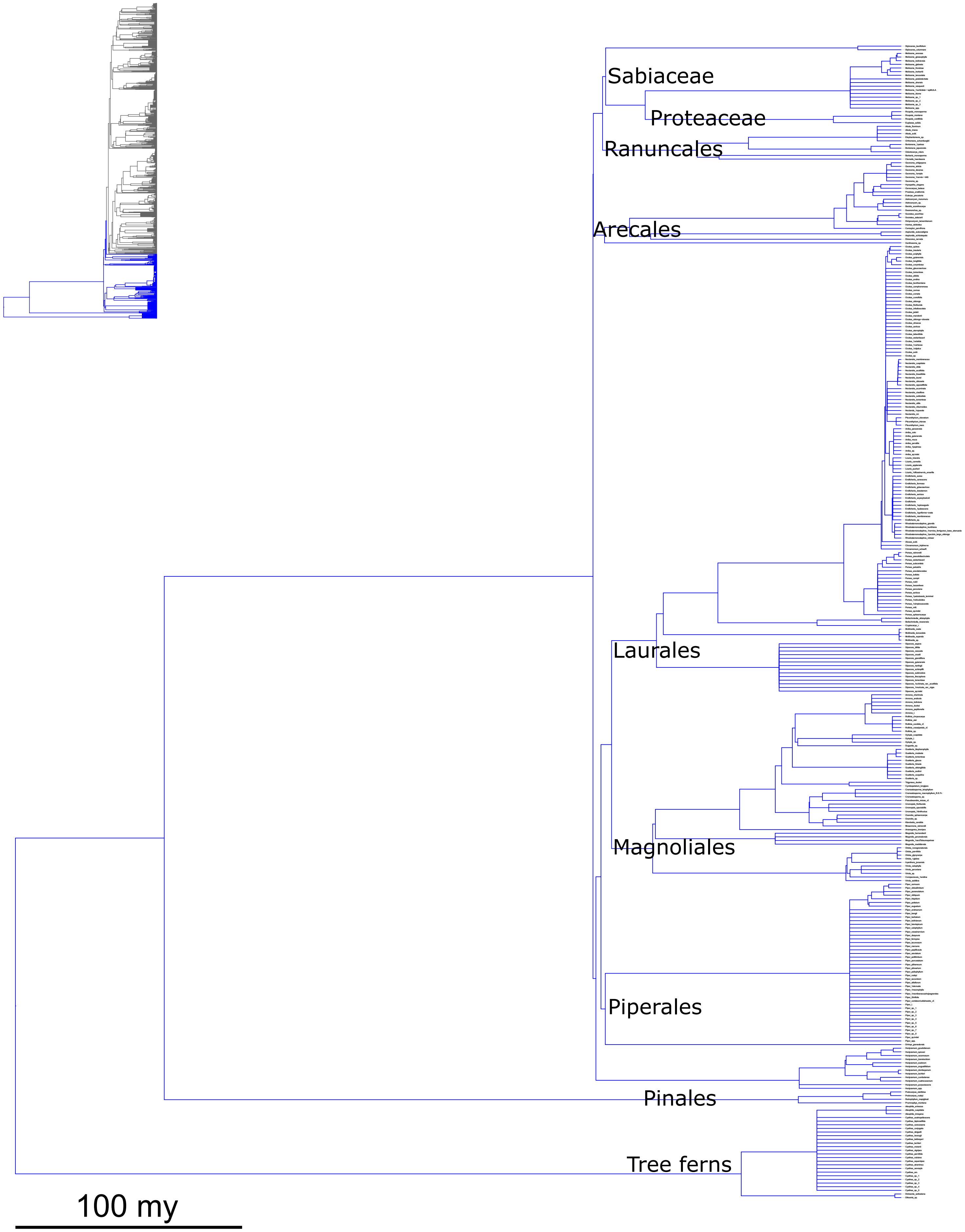
