## Appendix 2 for "Woody plant taxonomic, functional, and phylogenetic diversity decrease along elevational gradients in Andean tropical montane forests: environmental filtering and arrival of temperate taxa"

**Table A1. Information about the location, elevation, elevational belt, number of taxa, number and % of taxa over which functional and phylogenetic analysis were conducted, and taxonomic resolution of each plot of the four transects ranging elevational gradients.** Light grey cells indicate lack of functional data for Madidi and Pilón-Lajas transects.

| **Transect** | **Plot** | **Longitude (°)** | **Latitude (°)** | **Elev. (masl)** | **Belt** | ***n* taxa** | ***n* taxa with funct. data** | **% taxa with funct. data** | ***n* taxa with phyl. data** | **% taxa with phyl. data** | ***n* taxa (species level)** | **% taxa (species level)** | ***n* taxa (genera level)** | **% taxa (genera level)** | ***n* taxa (above genera level)** | **% taxa (above genera level)** |
| --- | --- | --- | --- | --- | --- | --- | --- | --- | --- | --- | --- | --- | --- | --- | --- | --- |
| PN Podocarpus -Ecuador- | Bombuscaro.A1 | -78,970967 | -4,119717 | 1084 | Premontane | 102 | 86 | 84 | 89 | 87 | 71 | 70 | 18 | 18 | 13 | 13 |
|  | Bombuscaro.A2 | -78,964472 | -4,11275 | 1053 |  | 66 | 58 | 88 | 60 | 91 | 45 | 68 | 15 | 23 | 6 | 9 |
|  | Bombuscaro.A3 | -78,968575 | -4,117603 | 1057 |  | 123 | 106 | 86 | 107 | 87 | 90 | 73 | 17 | 14 | 16 | 13 |
|  | Bombuscaro.A4 | -78,964869 | -4,115967 | 1104 |  | 134 | 116 | 87 | 114 | 85 | 95 | 71 | 19 | 14 | 20 | 15 |
|  | Bombuscaro.A5 | -78,961969 | -4,112369 | 1033 |  | 123 | 108 | 88 | 111 | 90 | 89 | 72 | 22 | 18 | 12 | 10 |
|  | Bombuscaro.A6 | -78,975627 | -4,121194 | 1084 |  | 79 | 72 | 91 | 68 | 86 | 52 | 66 | 16 | 20 | 11 | 14 |
|  | Bombuscaro.A7 | -78,978097 | -4,124908 | 1129 |  | 65 | 61 | 94 | 56 | 86 | 48 | 74 | 8 | 12 | 9 | 14 |
|  | Bombuscaro.A8 | -78,980158 | -4,127579 | 1216 |  | 61 | 50 | 82 | 58 | 95 | 45 | 74 | 13 | 21 | 3 | 5 |
|  | Bombuscaro.A9 | -78,980554 | -4,131513 | 1249 |  | 75 | 73 | 97 | 72 | 96 | 59 | 79 | 13 | 17 | 3 | 4 |
|  | Bombuscaro.A10 | -78,969242 | -4,114559 | 1130 |  | 104 | 95 | 91 | 96 | 92 | 84 | 81 | 12 | 12 | 8 | 8 |
|  | Bombuscaro.B1 | -79,009822 | -4,132517 | 1806 | Montane | 96 | 82 | 85 | 90 | 94 | 75 | 78 | 15 | 16 | 6 | 6 |
|  | Bombuscaro.B2 | -79,012364 | -4,131953 | 1837 |  | 64 | 51 | 80 | 53 | 83 | 45 | 70 | 8 | 13 | 11 | 17 |
|  | Bombuscaro.B3 | -79,0106 | -4,128777 | 1952 |  | 105 | 91 | 87 | 94 | 90 | 78 | 74 | 16 | 15 | 11 | 10 |
|  | Bombuscaro.B4 | -79,01511 | -4,132622 | 1851 |  | 99 | 75 | 76 | 92 | 93 | 78 | 79 | 14 | 14 | 7 | 7 |
|  | Bombuscaro.B5 | -79,016925 | -4,130683 | 1860 |  | 93 | 75 | 81 | 86 | 92 | 70 | 75 | 16 | 17 | 7 | 8 |
|  | Bombuscaro.B6 | -79,019635 | -4,130672 | 1901 |  | 84 | 72 | 86 | 78 | 93 | 61 | 73 | 17 | 20 | 6 | 7 |
|  | Bombuscaro.B7 | -79,013811 | -4,126534 | 2217 |  | 54 | 44 | 81 | 48 | 89 | 40 | 74 | 8 | 15 | 6 | 11 |
|  | Bombuscaro.B8 | -79,014515 | -4,129384 | 2060 |  | 77 | 68 | 88 | 74 | 96 | 64 | 83 | 10 | 13 | 3 | 4 |
|  | Bombuscaro.B9 | -79,022806 | -4,130336 | 2102 |  | 76 | 64 | 84 | 74 | 97 | 58 | 76 | 16 | 21 | 2 | 3 |
|  | Bombuscaro.B10 | -79,025295 | -4,128751 | 2190 |  | 60 | 53 | 88 | 57 | 95 | 46 | 77 | 11 | 18 | 3 | 5 |
|  | Bombuscaro.C1 | -79,024139 | -4,108305 | 2817 | Upper | 62 | 57 | 92 | 58 | 94 | 53 | 85 | 5 | 8 | 4 | 6 |
|  | Bombuscaro.C2 | -79,021398 | -4,106056 | 2796 |  | 51 | 45 | 88 | 50 | 98 | 45 | 88 | 5 | 10 | 1 | 2 |
|  | Bombuscaro.C3 | -79,021215 | -4,103533 | 2851 |  | 58 | 54 | 93 | 55 | 95 | 53 | 91 | 2 | 3 | 3 | 5 |
|  | Bombuscaro.C4 | -79,024935 | -4,103009 | 2906 |  | 50 | 48 | 96 | 50 | 100 | 47 | 94 | 3 | 6 | 0 | 0 |
|  | Bombuscaro.C5 | -79,025027 | -4,106183 | 2900 |  | 61 | 56 | 92 | 60 | 98 | 56 | 92 | 4 | 7 | 1 | 2 |
|  | Bombuscaro.C6 | -79,022835 | -4,111277 | 2729 |  | 47 | 38 | 81 | 43 | 91 | 39 | 83 | 4 | 9 | 4 | 9 |
|  | Bombuscaro.C7 | -79,02112 | -4,113053 | 2703 |  | 60 | 55 | 92 | 58 | 97 | 52 | 87 | 6 | 10 | 2 | 3 |
|  | Bombuscaro.C8 | -79,016544 | -4,102644 | 2738 |  | 55 | 35 | 64 | 41 | 75 | 36 | 65 | 5 | 9 | 14 | 25 |
|  | Bombuscaro.C9 | -79,018394 | -4,104557 | 2765 |  | 56 | 40 | 71 | 48 | 86 | 42 | 75 | 6 | 11 | 8 | 14 |
|  | Bombuscaro.C10 | -79,020317 | -4,117274 | 2674 |  | 32 | 25 | 78 | 29 | 91 | 27 | 84 | 2 | 6 | 3 | 9 |
| PN Río Abiseo -Peru- | Abiseo.A1 | -76,9006488 | -7,425792679 | 898 | Premontane | 82 | 77 | 94 | 64 | 78 | 36 | 44 | 28 | 34 | 18 | 22 |
|  | Abiseo.A2 | -76,8978395 | -7,425443058 | 819 |  | 50 | 48 | 96 | 39 | 78 | 27 | 54 | 12 | 24 | 11 | 22 |
|  | Abiseo.A3 | -76,8955746 | -7,42320153 | 941 |  | 102 | 91 | 89 | 82 | 80 | 40 | 39 | 42 | 41 | 20 | 20 |
|  | Abiseo.A4 | -76,8981523 | -7,422358751 | 963 |  | 104 | 91 | 88 | 83 | 80 | 43 | 41 | 40 | 38 | 21 | 20 |
|  | Abiseo.A5 | -76,9011234 | -7,422427413 | 1122 |  | 126 | 108 | 86 | 101 | 80 | 57 | 45 | 44 | 35 | 25 | 20 |
|  | Abiseo.A6 | -76,899327 | -7,428049534 | 780 |  | 118 | 100 | 85 | 99 | 84 | 55 | 47 | 44 | 37 | 19 | 16 |
|  | Abiseo.A7 | -76,9016754 | -7,426520599 | 912 |  | 106 | 89 | 84 | 85 | 80 | 41 | 39 | 44 | 42 | 21 | 20 |
|  | Abiseo.A8 | -76,9036926 | -7,428030831 | 793 |  | 108 | 93 | 86 | 96 | 89 | 55 | 51 | 41 | 38 | 12 | 11 |
|  | Abiseo.A9 | -76,8950303 | -7,42722709 | 745 |  | 101 | 91 | 90 | 80 | 79 | 40 | 40 | 40 | 40 | 21 | 21 |
|  | Abiseo.A10 | -76,9254655 | -7,452708757 | 841 |  | 99 | 87 | 88 | 84 | 85 | 44 | 44 | 40 | 40 | 15 | 15 |
|  | Abiseo.B1 | -77,379199 | -7,638242162 | 2117 | Montane | 68 | 66 | 97 | 37 | 54 | 18 | 26 | 19 | 28 | 31 | 46 |
|  | Abiseo.B2 | -77,3794573 | -7,640717188 | 2188 |  | 106 | 100 | 94 | 61 | 58 | 28 | 26 | 33 | 31 | 45 | 42 |
|  | Abiseo.B3 | -77,3767682 | -7,637695184 | 2152 |  | 92 | 87 | 95 | 53 | 58 | 27 | 29 | 26 | 28 | 39 | 42 |
|  | Abiseo.B4 | -77,3753508 | -7,633662924 | 2093 |  | 82 | 76 | 93 | 48 | 59 | 22 | 27 | 26 | 32 | 34 | 41 |
|  | Abiseo.B5 | -77,3737509 | -7,631014495 | 2144 |  | 64 | 62 | 97 | 35 | 55 | 18 | 28 | 17 | 27 | 29 | 45 |
|  | Abiseo.B6 | -77,3846728 | -7,641890475 | 2143 |  | 64 | 63 | 98 | 38 | 59 | 16 | 25 | 22 | 34 | 26 | 41 |
|  | Abiseo.B7 | -77,387777 | -7,64132199 | 2189 |  | 61 | 59 | 97 | 35 | 57 | 14 | 23 | 21 | 34 | 26 | 43 |
|  | Abiseo.B8 | -77,3898588 | -7,642639064 | 2233 |  | 50 | 45 | 90 | 33 | 66 | 13 | 26 | 20 | 40 | 17 | 34 |
|  | Abiseo.B9 | -77,3915448 | -7,644437335 | 2222 |  | 51 | 48 | 94 | 29 | 57 | 15 | 29 | 14 | 27 | 22 | 43 |
|  | Abiseo.B10 | -77,3946799 | -7,644555517 | 2101 |  | 46 | 44 | 96 | 26 | 57 | 13 | 28 | 13 | 28 | 20 | 43 |
|  | Abiseo.C1 | -77,4343626 | -7,662887997 | 2721 | Upper | 33 | 30 | 91 | 26 | 79 | 19 | 58 | 7 | 21 | 7 | 21 |
|  | Abiseo.C2 | -77,4332568 | -7,659604498 | 2767 |  | 30 | 29 | 97 | 17 | 57 | 13 | 43 | 4 | 13 | 13 | 43 |
|  | Abiseo.C3 | -77,4235593 | -7,660472591 | 2791 |  | 45 | 41 | 91 | 28 | 62 | 18 | 40 | 10 | 22 | 17 | 38 |
|  | Abiseo.C4 | -77,4379915 | -7,663807385 | 2759 |  | 36 | 33 | 92 | 23 | 64 | 16 | 44 | 7 | 19 | 13 | 36 |
|  | Abiseo.C5 | -77,4418907 | -7,664463109 | 2778 |  | 26 | 25 | 96 | 20 | 77 | 16 | 62 | 4 | 15 | 6 | 23 |
|  | Abiseo.C6 | -77,4471836 | -7,664857815 | 2866 |  | 47 | 43 | 91 | 29 | 62 | 22 | 47 | 7 | 15 | 18 | 38 |
|  | Abiseo.C7 | -77,4459941 | -7,667557811 | 2980 |  | 36 | 32 | 89 | 24 | 67 | 17 | 47 | 7 | 19 | 12 | 33 |
|  | Abiseo.C8 | -77,4322617 | -7,656645726 | 2775 |  | 54 | 49 | 91 | 35 | 65 | 24 | 44 | 11 | 20 | 19 | 35 |
|  | Abiseo.C9 | -77,4275339 | -7,660043499 | 2821 |  | 47 | 43 | 91 | 27 | 57 | 17 | 36 | 10 | 21 | 20 | 43 |
|  | Abiseo.C10 | -77,4301088 | -7,662008254 | 2810 |  | 43 | 37 | 86 | 31 | 72 | 22 | 51 | 9 | 21 | 12 | 28 |
| PN Madidi -Bolivia- | PT_Victop_372 | -68,3561217 | -15,4683937 | 1531 | Premontane | 82 |  |  | 81 | 99 | 69 | 84 | 12 | 15 | 1 | 1 |
|  | PT_Victop_373 | -68,3552565 | -15,4678488 | 1489 |  | 80 |  |  | 78 | 98 | 66 | 83 | 12 | 15 | 2 | 3 |
|  | PT_Victop_374 | -68,3497296 | -15,4615586 | 1493 |  | 101 |  |  | 101 | 100 | 83 | 82 | 18 | 18 | 0 | 0 |
|  | PT_Victop_375 | -68,3554672 | -15,4569648 | 1241 |  | 73 |  |  | 72 | 99 | 65 | 89 | 7 | 10 | 1 | 1 |
|  | PT_Victop_376 | -68,3563992 | -15,4600231 | 1224 |  | 77 |  |  | 76 | 99 | 63 | 82 | 13 | 17 | 1 | 1 |
|  | PT_Victop_377 | -68,3542573 | -15,4624036 | 1347 |  | 102 |  |  | 100 | 98 | 90 | 88 | 10 | 10 | 2 | 2 |
|  | PT_Victop_378 | -68,3530333 | -15,4603297 | 1386 |  | 83 |  |  | 82 | 99 | 66 | 80 | 16 | 19 | 1 | 1 |
|  | PT_Victop_379 | -68,3563142 | -15,4664325 | 1371 |  | 90 |  |  | 88 | 98 | 78 | 87 | 10 | 11 | 2 | 2 |
|  | PT_Victop_380 | -68,3554342 | -15,46464 | 1299 |  | 39 |  |  | 39 | 100 | 32 | 82 | 7 | 18 | 0 | 0 |
|  | PT_Lambra_434 | -68,3790793 | -15,6556119 | 2272 | Montane | 37 |  |  | 37 | 100 | 30 | 81 | 7 | 19 | 0 | 0 |
|  | PT_Lambra_435 | -68,3750348 | -15,657083 | 2019 |  | 75 |  |  | 73 | 97 | 62 | 83 | 11 | 15 | 2 | 3 |
|  | PT_Lambra_436 | -68,3742849 | -15,6490441 | 2251 |  | 36 |  |  | 35 | 97 | 30 | 83 | 5 | 14 | 1 | 3 |
|  | PT_Lambra_437 | -68,3703572 | -15,6489515 | 2179 |  | 56 |  |  | 55 | 98 | 43 | 77 | 12 | 21 | 1 | 2 |
|  | PT_Lambra_438 | -68,3707603 | -15,6513845 | 2045 |  | 58 |  |  | 57 | 98 | 46 | 79 | 11 | 19 | 1 | 2 |
|  | PT_Lambra_439 | -68,377215 | -15,6519453 | 2303 |  | 48 |  |  | 47 | 98 | 39 | 81 | 8 | 17 | 1 | 2 |
|  | PT_Lambra_440 | -68,3730615 | -15,6555496 | 2031 |  | 60 |  |  | 60 | 100 | 49 | 82 | 11 | 18 | 0 | 0 |
|  | PT_Lambra_441 | -68,3821549 | -15,6598334 | 2203 |  | 68 |  |  | 65 | 96 | 53 | 78 | 12 | 18 | 3 | 4 |
|  | PT_Lambra_442 | -68,3769472 | -15,6541233 | 2210 |  | 68 |  |  | 67 | 99 | 53 | 78 | 14 | 21 | 1 | 1 |
|  | PT_Cocapu_396 | -68,3933759 | -15,5531439 | 2801 | Upper | 36 |  |  | 36 | 100 | 29 | 81 | 7 | 19 | 0 | 0 |
|  | PT_Cocapu_397 | -68,3931547 | -15,5554305 | 2833 |  | 35 |  |  | 35 | 100 | 30 | 86 | 5 | 14 | 0 | 0 |
|  | PT_Cocapu_398 | -68,3952322 | -15,5561504 | 2888 |  | 36 |  |  | 36 | 100 | 32 | 89 | 4 | 11 | 0 | 0 |
|  | PT_Cocapu_399 | -68,3943118 | -15,5583718 | 2985 |  | 45 |  |  | 45 | 100 | 36 | 80 | 9 | 20 | 0 | 0 |
|  | PT_Cocapu_400 | -68,400043 | -15,5598249 | 3021 |  | 43 |  |  | 43 | 100 | 37 | 86 | 6 | 14 | 0 | 0 |
|  | PT_Cocapu_401 | -68,4022458 | -15,5624165 | 3106 |  | 38 |  |  | 38 | 100 | 34 | 89 | 4 | 11 | 0 | 0 |
|  | PT_Cocapu_402 | -68,3998441 | -15,5608911 | 3103 |  | 32 |  |  | 32 | 100 | 28 | 88 | 4 | 13 | 0 | 0 |
|  | PT_Cocapu_403 | -68,3952969 | -15,562732 | 3075 |  | 42 |  |  | 42 | 100 | 35 | 83 | 7 | 17 | 0 | 0 |
|  | PT_Cocapu_404 | -68,397189 | -15,5599075 | 3087 |  | 40 |  |  | 40 | 100 | 34 | 85 | 6 | 15 | 0 | 0 |
| Pilón-Lajas -Bolivia- | PT_Culi_353 | -68,8308116 | -14,7259315 | 1166 | Premontane | 63 |  |  | 60 | 95 | 52 | 83 | 8 | 13 | 3 | 5 |
|  | PT_Culi_354 | -68,8373217 | -14,7288023 | 1239 |  | 72 |  |  | 72 | 100 | 66 | 92 | 6 | 8 | 0 | 0 |
|  | PT_Culi_355 | -68,8444751 | -14,7287439 | 1496 |  | 100 |  |  | 98 | 98 | 85 | 85 | 13 | 13 | 2 | 2 |
|  | PT_Culi_356 | -68,8441387 | -14,7314652 | 1367 |  | 75 |  |  | 74 | 99 | 66 | 88 | 8 | 11 | 1 | 1 |
|  | PT_Culi_357 | -68,8545159 | -14,745251 | 1447 |  | 79 |  |  | 78 | 99 | 68 | 86 | 10 | 13 | 1 | 1 |
|  | PT_Culi_358 | -68,8528823 | -14,7429173 | 1330 |  | 89 |  |  | 88 | 99 | 80 | 90 | 8 | 9 | 1 | 1 |
|  | PT_Culi_359 | -68,8508043 | -14,7384314 | 1254 |  | 67 |  |  | 66 | 99 | 57 | 85 | 9 | 13 | 1 | 1 |
|  | PT_Culi_360 | -68,8493033 | -14,7325626 | 1469 |  | 91 |  |  | 90 | 99 | 80 | 88 | 10 | 11 | 1 | 1 |
|  | PT_Culi_361 | -68,8474342 | -14,7351382 | 1341 |  | 72 |  |  | 72 | 100 | 64 | 89 | 8 | 11 | 0 | 0 |
|  | PT_Santaa_344 | -68,9715761 | -14,7716869 | 2199 | Montane | 37 |  |  | 37 | 100 | 32 | 86 | 5 | 14 | 0 | 0 |
|  | PT_Santaa_345 | -68,974791 | -14,7727813 | 2281 |  | 31 |  |  | 30 | 97 | 25 | 81 | 5 | 16 | 1 | 3 |
|  | PT_Santaa_346 | -68,9609207 | -14,7582679 | 2252 |  | 41 |  |  | 41 | 100 | 37 | 90 | 4 | 10 | 0 | 0 |
|  | PT_Santaa_347 | -68,9794838 | -14,768686 | 2169 |  | 34 |  |  | 34 | 100 | 28 | 82 | 6 | 18 | 0 | 0 |
|  | PT_Santaa_348 | -68,9707031 | -14,7681426 | 2139 |  | 48 |  |  | 48 | 100 | 40 | 83 | 8 | 17 | 0 | 0 |
|  | PT_Santaa_349 | -68,9634106 | -14,7591363 | 2196 |  | 41 |  |  | 41 | 100 | 33 | 80 | 8 | 20 | 0 | 0 |
|  | PT_Santaa_350 | -68,9643766 | -14,7612521 | 2228 |  | 41 |  |  | 41 | 100 | 35 | 85 | 6 | 15 | 0 | 0 |
|  | PT_Santaa_351 | -68,9661696 | -14,7625725 | 2244 |  | 29 |  |  | 29 | 100 | 28 | 97 | 1 | 3 | 0 | 0 |
|  | PT_Santaa_352 | -68,9817786 | -14,7715704 | 2223 |  | 47 |  |  | 47 | 100 | 42 | 89 | 5 | 11 | 0 | 0 |
|  | PT_Piara_387 | -69,0150067 | -14,7766158 | 2821 | Upper | 34 |  |  | 33 | 97 | 26 | 76 | 7 | 21 | 1 | 3 |
|  | PT_Piara_388 | -69,0155923 | -14,7791473 | 2812 |  | 37 |  |  | 37 | 100 | 32 | 86 | 5 | 14 | 0 | 0 |
|  | PT_Piara_389 | -69,014905 | -14,7837585 | 2789 |  | 30 |  |  | 30 | 100 | 24 | 80 | 6 | 20 | 0 | 0 |
|  | PT_Piara_390 | -69,0204339 | -14,7827635 | 2826 |  | 28 |  |  | 28 | 100 | 23 | 82 | 5 | 18 | 0 | 0 |
|  | PT_Piara_391 | -69,0213261 | -14,7835862 | 2802 |  | 33 |  |  | 32 | 97 | 26 | 79 | 6 | 18 | 1 | 3 |
|  | PT_Piara_392 | -69,0265767 | -14,7869671 | 3061 |  | 40 |  |  | 39 | 98 | 30 | 75 | 9 | 23 | 1 | 3 |
|  | PT_Piara_393 | -69,0194122 | -14,7879172 | 2955 |  | 38 |  |  | 36 | 95 | 31 | 82 | 5 | 13 | 2 | 5 |
|  | PT_Piara_394 | -69,0299232 | -14,79599 | 2987 |  | 44 |  |  | 43 | 98 | 37 | 84 | 6 | 14 | 1 | 2 |
|  | PT_Piara_395 | -69,0257596 | -14,7927899 | 2975 |  | 28 |  |  | 27 | 96 | 23 | 82 | 4 | 14 | 1 | 4 |

**Table A2. Bioclimatic variables values of each plot of the four transects ranging elevational gradients.** Data obtained from CHELSA climatological dataset (Karger et al. 2017). Temp.: temperature, Seas.: seasonality, Quart.: quarter, Prec.: precipitation, Month.: monthly.

| **Transect** | **Plot** | **Mean Ann. Temp. (°C)** | **Mean Diurnal Temp. Range (°C)** | **Iso-ther-ma-lity (°C)** | **Temp. Seas. (°C)** | **Mean Daily Max. Temp. Warmest Month (°C)** | **Mean Daily Max. Temp. Coldest Month (°C)** | **Ann. Temp. Range (°C)** | **Mean Daily Temp. Wettest Quart. (°C)** | **Mean Daily Temp. Driest Quart. (°C)** | **Mean Daily Temp. Warmest Quart. (°C)** | **Mean Daily Temp. Coldest Quart. (°C)** | **Ann. Prec. (kg/ m^2^)** | **Prec. Wettest Month (kg/ m^2^)** | **Prec. Driest Month (kg/ m^2^)** | **Prec. Seas. (kg/ m^2^)** | **Mean Month. Precip. Wettest Quart. (kg/ m^2^)** | **Mean Month. Precip. Driest Quart. (kg/ m^2^)** | **Mean Month. Precip. Warmest Quart. (kg/ m^2^)** | **Mean Month. Precip. Coldest Quart. (kg/ m^2^)** |
| --- | --- | --- | --- | --- | --- | --- | --- | --- | --- | --- | --- | --- | --- | --- | --- | --- | --- | --- | --- | --- |
| PN Podocarpus -Ecuador- | Bombuscaro.A1 | 209 | 75 | 646 | 609 | 270 | 153 | 117 | 210 | 202 | 217 | 198 | 1046 | 145 | 65 | 25 | 398 | 203 | 240 | 227 |
|  | Bombuscaro.A2 | 211 | 75 | 645 | 615 | 272 | 155 | 117 | 212 | 204 | 219 | 200 | 1046 | 143 | 65 | 24 | 393 | 203 | 238 | 228 |
|  | Bombuscaro.A3 | 209 | 75 | 646 | 609 | 270 | 153 | 117 | 210 | 202 | 217 | 198 | 1046 | 145 | 65 | 25 | 398 | 203 | 240 | 227 |
|  | Bombuscaro.A4 | 211 | 75 | 645 | 615 | 272 | 155 | 117 | 212 | 204 | 219 | 200 | 1046 | 143 | 65 | 24 | 393 | 203 | 238 | 228 |
|  | Bombuscaro.A5 | 211 | 75 | 645 | 615 | 272 | 155 | 117 | 212 | 204 | 219 | 200 | 1046 | 143 | 65 | 24 | 393 | 203 | 238 | 228 |
|  | Bombuscaro.A6 | 199 | 76 | 649 | 595 | 259 | 143 | 116 | 199 | 191 | 206 | 187 | 1111 | 157 | 65 | 27 | 429 | 215 | 257 | 241 |
|  | Bombuscaro.A7 | 199 | 76 | 649 | 595 | 259 | 143 | 116 | 199 | 191 | 206 | 187 | 1111 | 157 | 65 | 27 | 429 | 215 | 257 | 241 |
|  | Bombuscaro.A8 | 207 | 76 | 646 | 604 | 268 | 151 | 117 | 208 | 200 | 215 | 196 | 957 | 135 | 58 | 26 | 370 | 182 | 217 | 206 |
|  | Bombuscaro.A9 | 207 | 76 | 646 | 604 | 268 | 151 | 117 | 208 | 200 | 215 | 196 | 957 | 135 | 58 | 26 | 370 | 182 | 217 | 206 |
|  | Bombuscaro.A10 | 201 | 75 | 648 | 598 | 262 | 145 | 117 | 202 | 194 | 209 | 190 | 1116 | 155 | 67 | 25 | 424 | 218 | 258 | 244 |
|  | Bombuscaro.B1 | 165 | 76 | 658 | 544 | 225 | 110 | 115 | 166 | 169 | 172 | 155 | 1390 | 208 | 68 | 31 | 561 | 244 | 319 | 304 |
|  | Bombuscaro.B2 | 165 | 76 | 658 | 544 | 225 | 110 | 115 | 166 | 169 | 172 | 155 | 1390 | 208 | 68 | 31 | 561 | 244 | 319 | 304 |
|  | Bombuscaro.B3 | 165 | 76 | 658 | 544 | 225 | 110 | 115 | 166 | 169 | 172 | 155 | 1390 | 208 | 68 | 31 | 561 | 244 | 319 | 304 |
|  | Bombuscaro.B4 | 165 | 76 | 658 | 544 | 225 | 110 | 115 | 166 | 169 | 172 | 155 | 1390 | 208 | 68 | 31 | 561 | 244 | 319 | 304 |
|  | Bombuscaro.B5 | 164 | 76 | 660 | 537 | 224 | 108 | 115 | 165 | 170 | 171 | 154 | 1391 | 210 | 66 | 32 | 565 | 240 | 315 | 305 |
|  | Bombuscaro.B6 | 164 | 76 | 660 | 537 | 224 | 108 | 115 | 165 | 170 | 171 | 154 | 1391 | 210 | 66 | 32 | 565 | 240 | 315 | 305 |
|  | Bombuscaro.B7 | 165 | 76 | 658 | 544 | 225 | 110 | 115 | 166 | 169 | 172 | 155 | 1390 | 208 | 68 | 31 | 561 | 244 | 319 | 304 |
|  | Bombuscaro.B8 | 165 | 76 | 658 | 544 | 225 | 110 | 115 | 166 | 169 | 172 | 155 | 1390 | 208 | 68 | 31 | 561 | 244 | 319 | 304 |
|  | Bombuscaro.B9 | 164 | 76 | 660 | 537 | 224 | 108 | 115 | 165 | 170 | 171 | 154 | 1391 | 210 | 66 | 32 | 565 | 240 | 315 | 305 |
|  | Bombuscaro.B10 | 163 | 76 | 660 | 533 | 222 | 107 | 115 | 163 | 169 | 170 | 152 | 1339 | 204 | 62 | 33 | 547 | 226 | 299 | 296 |
|  | Bombuscaro.C1 | 122 | 76 | 667 | 495 | 180 | 66 | 114 | 123 | 127 | 127 | 112 | 1614 | 236 | 89 | 29 | 641 | 302 | 360 | 349 |
|  | Bombuscaro.C2 | 122 | 76 | 667 | 495 | 180 | 66 | 114 | 123 | 127 | 127 | 112 | 1614 | 236 | 89 | 29 | 641 | 302 | 360 | 349 |
|  | Bombuscaro.C3 | 122 | 76 | 667 | 495 | 180 | 66 | 114 | 123 | 127 | 127 | 112 | 1614 | 236 | 89 | 29 | 641 | 302 | 360 | 349 |
|  | Bombuscaro.C4 | 122 | 76 | 667 | 495 | 180 | 66 | 114 | 123 | 127 | 127 | 112 | 1614 | 236 | 89 | 29 | 641 | 302 | 360 | 349 |
|  | Bombuscaro.C5 | 122 | 76 | 667 | 495 | 180 | 66 | 114 | 123 | 127 | 127 | 112 | 1614 | 236 | 89 | 29 | 641 | 302 | 360 | 349 |
|  | Bombuscaro.C6 | 131 | 76 | 667 | 502 | 190 | 76 | 114 | 132 | 136 | 137 | 121 | 1544 | 229 | 78 | 31 | 619 | 276 | 349 | 337 |
|  | Bombuscaro.C7 | 131 | 76 | 667 | 502 | 190 | 76 | 114 | 132 | 136 | 137 | 121 | 1544 | 229 | 78 | 31 | 619 | 276 | 349 | 337 |
|  | Bombuscaro.C8 | 138 | 76 | 665 | 513 | 197 | 82 | 114 | 139 | 143 | 144 | 128 | 1610 | 234 | 86 | 29 | 635 | 297 | 364 | 352 |
|  | Bombuscaro.C9 | 122 | 76 | 667 | 495 | 180 | 66 | 114 | 123 | 127 | 127 | 112 | 1614 | 236 | 89 | 29 | 641 | 302 | 360 | 349 |
|  | Bombuscaro.C10 | 141 | 76 | 665 | 514 | 200 | 85 | 114 | 142 | 144 | 147 | 131 | 1467 | 220 | 71 | 32 | 593 | 255 | 332 | 323 |
| PN Río Abiseo -Peru- | Abiseo.A1 | 227 | 74 | 646 | 592 | 280 | 166 | 114 | 228 | 219 | 233 | 215 | 1802 | 224 | 80 | 32 | 670 | 242 | 560 | 262 |
|  | Abiseo.A2 | 237 | 74 | 644 | 599 | 290 | 176 | 114 | 238 | 229 | 243 | 226 | 1618 | 201 | 71 | 32 | 599 | 214 | 468 | 231 |
|  | Abiseo.A3 | 235 | 74 | 645 | 595 | 288 | 174 | 114 | 236 | 227 | 241 | 223 | 1867 | 229 | 80 | 32 | 681 | 242 | 594 | 263 |
|  | Abiseo.A4 | 235 | 74 | 645 | 595 | 288 | 174 | 114 | 236 | 227 | 241 | 223 | 1867 | 229 | 80 | 32 | 681 | 242 | 594 | 263 |
|  | Abiseo.A5 | 221 | 74 | 646 | 587 | 274 | 160 | 114 | 223 | 213 | 228 | 210 | 2007 | 245 | 86 | 32 | 729 | 260 | 638 | 282 |
|  | Abiseo.A6 | 237 | 74 | 644 | 599 | 290 | 176 | 114 | 238 | 229 | 243 | 226 | 1618 | 201 | 71 | 32 | 599 | 214 | 468 | 231 |
|  | Abiseo.A7 | 227 | 74 | 646 | 592 | 280 | 166 | 114 | 228 | 219 | 233 | 215 | 1802 | 224 | 80 | 32 | 670 | 242 | 560 | 262 |
|  | Abiseo.A8 | 227 | 74 | 646 | 592 | 280 | 166 | 114 | 228 | 219 | 233 | 215 | 1802 | 224 | 80 | 32 | 670 | 242 | 560 | 262 |
|  | Abiseo.A9 | 237 | 74 | 644 | 599 | 290 | 176 | 114 | 238 | 229 | 243 | 226 | 1618 | 201 | 71 | 32 | 599 | 214 | 468 | 231 |
|  | Abiseo.A10 | 239 | 74 | 644 | 601 | 292 | 178 | 115 | 240 | 231 | 245 | 228 | 1563 | 195 | 66 | 33 | 575 | 200 | 449 | 212 |
|  | Abiseo.B1 | 161 | 78 | 656 | 496 | 216 | 97 | 119 | 162 | 151 | 166 | 151 | 1058 | 169 | 30 | 48 | 469 | 96 | 298 | 96 |
|  | Abiseo.B2 | 161 | 78 | 656 | 496 | 216 | 97 | 119 | 162 | 151 | 166 | 151 | 1058 | 169 | 30 | 48 | 469 | 96 | 298 | 96 |
|  | Abiseo.B3 | 161 | 78 | 656 | 496 | 216 | 97 | 119 | 162 | 151 | 166 | 151 | 1058 | 169 | 30 | 48 | 469 | 96 | 298 | 96 |
|  | Abiseo.B4 | 161 | 78 | 656 | 496 | 216 | 97 | 119 | 162 | 151 | 166 | 151 | 1058 | 169 | 30 | 48 | 469 | 96 | 298 | 96 |
|  | Abiseo.B5 | 166 | 78 | 656 | 500 | 221 | 102 | 119 | 167 | 156 | 171 | 156 | 1079 | 170 | 33 | 46 | 470 | 105 | 301 | 105 |
|  | Abiseo.B6 | 152 | 78 | 658 | 489 | 207 | 89 | 119 | 153 | 142 | 157 | 142 | 1019 | 163 | 27 | 49 | 455 | 87 | 287 | 87 |
|  | Abiseo.B7 | 147 | 78 | 660 | 485 | 201 | 83 | 118 | 148 | 137 | 151 | 137 | 1104 | 177 | 34 | 47 | 487 | 108 | 309 | 108 |
|  | Abiseo.B8 | 152 | 78 | 658 | 489 | 207 | 89 | 119 | 153 | 142 | 157 | 142 | 1019 | 163 | 27 | 49 | 455 | 87 | 287 | 87 |
|  | Abiseo.B9 | 152 | 78 | 658 | 489 | 207 | 89 | 119 | 153 | 142 | 157 | 142 | 1019 | 163 | 27 | 49 | 455 | 87 | 287 | 87 |
|  | Abiseo.B10 | 153 | 78 | 657 | 488 | 208 | 89 | 119 | 154 | 143 | 158 | 143 | 1080 | 174 | 31 | 48 | 482 | 99 | 303 | 99 |
|  | Abiseo.C1 | 128 | 78 | 662 | 464 | 183 | 64 | 119 | 129 | 118 | 132 | 118 | 1372 | 220 | 37 | 50 | 617 | 118 | 373 | 118 |
|  | Abiseo.C2 | 128 | 78 | 662 | 465 | 182 | 64 | 118 | 129 | 118 | 132 | 118 | 1375 | 219 | 36 | 50 | 617 | 115 | 372 | 115 |
|  | Abiseo.C3 | 129 | 78 | 661 | 470 | 184 | 66 | 118 | 130 | 120 | 134 | 120 | 1393 | 220 | 36 | 50 | 621 | 115 | 376 | 115 |
|  | Abiseo.C4 | 128 | 78 | 662 | 464 | 183 | 64 | 119 | 129 | 118 | 132 | 118 | 1372 | 220 | 37 | 50 | 617 | 118 | 373 | 118 |
|  | Abiseo.C5 | 124 | 79 | 662 | 461 | 179 | 60 | 119 | 125 | 117 | 128 | 114 | 1340 | 217 | 37 | 50 | 605 | 117 | 366 | 118 |
|  | Abiseo.C6 | 124 | 79 | 662 | 461 | 179 | 60 | 119 | 125 | 117 | 128 | 114 | 1340 | 217 | 37 | 50 | 605 | 117 | 366 | 118 |
|  | Abiseo.C7 | 116 | 78 | 663 | 456 | 170 | 52 | 118 | 117 | 109 | 120 | 106 | 1418 | 227 | 37 | 51 | 638 | 117 | 385 | 118 |
|  | Abiseo.C8 | 135 | 78 | 661 | 470 | 190 | 72 | 119 | 137 | 126 | 140 | 126 | 1297 | 207 | 36 | 49 | 579 | 115 | 352 | 115 |
|  | Abiseo.C9 | 128 | 78 | 662 | 465 | 182 | 64 | 118 | 129 | 118 | 132 | 118 | 1375 | 219 | 36 | 50 | 617 | 115 | 372 | 115 |
|  | Abiseo.C10 | 128 | 78 | 662 | 465 | 182 | 64 | 118 | 129 | 118 | 132 | 118 | 1375 | 219 | 36 | 50 | 617 | 115 | 372 | 115 |
| PN Madidi -Bolivia- | PT_Victop_372 | 188 | 71 | 566 | 1465 | 238 | 113 | 125 | 201 | 162 | 203 | 162 | 1265 | 216 | 19 | 62 | 611 | 61 | 482 | 61 |
|  | PT_Victop_373 | 188 | 71 | 566 | 1465 | 238 | 113 | 125 | 201 | 162 | 203 | 162 | 1265 | 216 | 19 | 62 | 611 | 61 | 482 | 61 |
|  | PT_Victop_374 | 194 | 71 | 566 | 1458 | 245 | 120 | 125 | 208 | 169 | 209 | 169 | 1204 | 202 | 19 | 61 | 575 | 61 | 461 | 61 |
|  | PT_Victop_375 | 210 | 71 | 566 | 1449 | 260 | 136 | 125 | 223 | 184 | 225 | 184 | 1043 | 183 | 15 | 65 | 514 | 47 | 398 | 47 |
|  | PT_Victop_376 | 200 | 71 | 566 | 1456 | 250 | 125 | 125 | 213 | 174 | 215 | 174 | 1181 | 205 | 17 | 64 | 578 | 55 | 452 | 55 |
|  | PT_Victop_377 | 200 | 71 | 566 | 1456 | 250 | 125 | 125 | 213 | 174 | 215 | 174 | 1181 | 205 | 17 | 64 | 578 | 55 | 452 | 55 |
|  | PT_Victop_378 | 200 | 71 | 566 | 1456 | 250 | 125 | 125 | 213 | 174 | 215 | 174 | 1181 | 205 | 17 | 64 | 578 | 55 | 452 | 55 |
|  | PT_Victop_379 | 200 | 71 | 566 | 1456 | 250 | 125 | 125 | 213 | 174 | 215 | 174 | 1181 | 205 | 17 | 64 | 578 | 55 | 452 | 55 |
|  | PT_Victop_380 | 200 | 71 | 566 | 1456 | 250 | 125 | 125 | 213 | 174 | 215 | 174 | 1181 | 205 | 17 | 64 | 578 | 55 | 452 | 55 |
|  | PT_Lambra_434 | 151 | 73 | 567 | 1516 | 203 | 74 | 130 | 165 | 125 | 167 | 125 | 806 | 144 | 12 | 65 | 403 | 40 | 306 | 40 |
|  | PT_Lambra_435 | 170 | 73 | 567 | 1506 | 222 | 93 | 129 | 184 | 144 | 186 | 144 | 772 | 137 | 12 | 64 | 385 | 38 | 295 | 38 |
|  | PT_Lambra_436 | 148 | 73 | 567 | 1514 | 200 | 71 | 129 | 162 | 122 | 164 | 122 | 884 | 157 | 13 | 65 | 441 | 43 | 338 | 43 |
|  | PT_Lambra_437 | 148 | 73 | 567 | 1514 | 200 | 71 | 129 | 162 | 122 | 164 | 122 | 884 | 157 | 13 | 65 | 441 | 43 | 338 | 43 |
|  | PT_Lambra_438 | 170 | 73 | 567 | 1506 | 222 | 93 | 129 | 184 | 144 | 186 | 144 | 772 | 137 | 12 | 64 | 385 | 38 | 295 | 38 |
|  | PT_Lambra_439 | 151 | 73 | 567 | 1516 | 203 | 74 | 130 | 165 | 125 | 167 | 125 | 806 | 144 | 12 | 65 | 403 | 40 | 306 | 40 |
|  | PT_Lambra_440 | 170 | 73 | 567 | 1506 | 222 | 93 | 129 | 184 | 144 | 186 | 144 | 772 | 137 | 12 | 64 | 385 | 38 | 295 | 38 |
|  | PT_Lambra_441 | 168 | 74 | 566 | 1510 | 220 | 90 | 130 | 182 | 142 | 184 | 142 | 804 | 145 | 12 | 66 | 406 | 38 | 308 | 38 |
|  | PT_Lambra_442 | 151 | 73 | 567 | 1516 | 203 | 74 | 130 | 165 | 125 | 167 | 125 | 806 | 144 | 12 | 65 | 403 | 40 | 306 | 40 |
|  | PT_Cocapu_396 | 129 | 73 | 569 | 1509 | 180 | 52 | 128 | 143 | 103 | 144 | 103 | 1567 | 271 | 26 | 62 | 761 | 82 | 587 | 82 |
|  | PT_Cocapu_397 | 129 | 73 | 569 | 1509 | 180 | 52 | 128 | 143 | 103 | 144 | 103 | 1567 | 271 | 26 | 62 | 761 | 82 | 587 | 82 |
|  | PT_Cocapu_398 | 129 | 73 | 569 | 1509 | 180 | 52 | 128 | 143 | 103 | 144 | 103 | 1567 | 271 | 26 | 62 | 761 | 82 | 587 | 82 |
|  | PT_Cocapu_399 | 129 | 73 | 569 | 1509 | 180 | 52 | 128 | 143 | 103 | 144 | 103 | 1567 | 271 | 26 | 62 | 761 | 82 | 587 | 82 |
|  | PT_Cocapu_400 | 112 | 73 | 569 | 1522 | 163 | 36 | 128 | 127 | 86 | 128 | 86 | 1608 | 279 | 27 | 62 | 783 | 85 | 602 | 85 |
|  | PT_Cocapu_401 | 114 | 73 | 569 | 1523 | 165 | 37 | 128 | 129 | 88 | 129 | 88 | 1605 | 279 | 26 | 62 | 783 | 84 | 602 | 84 |
|  | PT_Cocapu_402 | 112 | 73 | 569 | 1522 | 163 | 36 | 128 | 127 | 86 | 128 | 86 | 1608 | 279 | 27 | 62 | 783 | 85 | 602 | 85 |
|  | PT_Cocapu_403 | 112 | 73 | 569 | 1522 | 163 | 36 | 128 | 127 | 86 | 128 | 86 | 1608 | 279 | 27 | 62 | 783 | 85 | 602 | 85 |
|  | PT_Cocapu_404 | 112 | 73 | 569 | 1522 | 163 | 36 | 128 | 127 | 86 | 128 | 86 | 1608 | 279 | 27 | 62 | 783 | 85 | 602 | 85 |
| Pilón-Lajas -Bolivia- | PT_Culi_353 | 199 | 71 | 586 | 1341 | 249 | 127 | 121 | 210 | 176 | 213 | 176 | 747 | 117 | 10 | 62 | 345 | 32 | 284 | 32 |
|  | PT_Culi_354 | 197 | 71 | 586 | 1337 | 246 | 125 | 121 | 207 | 173 | 211 | 173 | 699 | 110 | 9 | 62 | 324 | 31 | 265 | 31 |
|  | PT_Culi_355 | 189 | 71 | 587 | 1339 | 238 | 117 | 121 | 199 | 165 | 203 | 165 | 745 | 117 | 10 | 62 | 347 | 32 | 281 | 32 |
|  | PT_Culi_356 | 189 | 71 | 587 | 1339 | 238 | 117 | 121 | 199 | 165 | 203 | 165 | 745 | 117 | 10 | 62 | 347 | 32 | 281 | 32 |
|  | PT_Culi_357 | 188 | 72 | 587 | 1343 | 238 | 116 | 122 | 199 | 165 | 202 | 165 | 821 | 131 | 10 | 64 | 391 | 34 | 308 | 34 |
|  | PT_Culi_358 | 188 | 72 | 587 | 1343 | 238 | 116 | 122 | 199 | 165 | 202 | 165 | 821 | 131 | 10 | 64 | 391 | 34 | 308 | 34 |
|  | PT_Culi_359 | 185 | 72 | 588 | 1342 | 235 | 113 | 122 | 196 | 162 | 199 | 162 | 767 | 120 | 10 | 63 | 358 | 32 | 288 | 32 |
|  | PT_Culi_360 | 189 | 71 | 587 | 1339 | 238 | 117 | 121 | 199 | 165 | 203 | 165 | 745 | 117 | 10 | 62 | 347 | 32 | 281 | 32 |
|  | PT_Culi_361 | 188 | 71 | 587 | 1343 | 238 | 116 | 122 | 199 | 165 | 202 | 165 | 738 | 116 | 9 | 63 | 344 | 31 | 278 | 31 |
|  | PT_Santaa_344 | 152 | 74 | 590 | 1365 | 203 | 78 | 125 | 163 | 128 | 166 | 128 | 848 | 138 | 9 | 65 | 411 | 31 | 318 | 31 |
|  | PT_Santaa_345 | 152 | 74 | 590 | 1365 | 203 | 78 | 125 | 163 | 128 | 166 | 128 | 848 | 138 | 9 | 65 | 411 | 31 | 318 | 31 |
|  | PT_Santaa_346 | 159 | 73 | 590 | 1355 | 209 | 85 | 124 | 169 | 135 | 172 | 135 | 754 | 120 | 8 | 64 | 359 | 30 | 283 | 30 |
|  | PT_Santaa_347 | 149 | 74 | 591 | 1361 | 199 | 74 | 125 | 160 | 125 | 163 | 125 | 770 | 124 | 8 | 65 | 371 | 30 | 288 | 30 |
|  | PT_Santaa_348 | 152 | 74 | 590 | 1365 | 203 | 78 | 125 | 163 | 128 | 166 | 128 | 848 | 138 | 9 | 65 | 411 | 31 | 318 | 31 |
|  | PT_Santaa_349 | 142 | 73 | 591 | 1365 | 192 | 68 | 124 | 153 | 118 | 156 | 118 | 858 | 139 | 9 | 65 | 415 | 31 | 321 | 31 |
|  | PT_Santaa_350 | 142 | 73 | 591 | 1365 | 192 | 68 | 124 | 153 | 118 | 156 | 118 | 858 | 139 | 9 | 65 | 415 | 31 | 321 | 31 |
|  | PT_Santaa_351 | 142 | 73 | 591 | 1365 | 192 | 68 | 124 | 153 | 118 | 156 | 118 | 858 | 139 | 9 | 65 | 415 | 31 | 321 | 31 |
|  | PT_Santaa_352 | 149 | 74 | 591 | 1361 | 199 | 74 | 125 | 160 | 125 | 163 | 125 | 770 | 124 | 8 | 65 | 371 | 30 | 288 | 30 |
|  | PT_Piara_387 | 128 | 74 | 592 | 1375 | 179 | 53 | 126 | 140 | 104 | 142 | 104 | 833 | 135 | 8 | 65 | 403 | 30 | 314 | 30 |
|  | PT_Piara_388 | 128 | 74 | 592 | 1375 | 179 | 53 | 126 | 140 | 104 | 142 | 104 | 833 | 135 | 8 | 65 | 403 | 30 | 314 | 30 |
|  | PT_Piara_389 | 106 | 74 | 592 | 1389 | 157 | 31 | 126 | 118 | 82 | 120 | 82 | 1070 | 175 | 10 | 66 | 523 | 36 | 405 | 36 |
|  | PT_Piara_390 | 114 | 74 | 592 | 1380 | 165 | 39 | 126 | 126 | 90 | 128 | 90 | 818 | 132 | 8 | 65 | 395 | 30 | 308 | 30 |
|  | PT_Piara_391 | 116 | 75 | 592 | 1382 | 166 | 40 | 126 | 127 | 92 | 130 | 92 | 926 | 153 | 8 | 67 | 456 | 32 | 348 | 32 |
|  | PT_Piara_392 | 101 | 75 | 593 | 1389 | 152 | 26 | 126 | 113 | 77 | 115 | 77 | 910 | 152 | 8 | 68 | 452 | 30 | 340 | 30 |
|  | PT_Piara_393 | 116 | 75 | 592 | 1382 | 166 | 40 | 126 | 127 | 92 | 130 | 92 | 926 | 153 | 8 | 67 | 456 | 32 | 348 | 32 |
|  | PT_Piara_394 | 109 | 75 | 592 | 1389 | 160 | 34 | 126 | 121 | 85 | 123 | 85 | 1032 | 174 | 9 | 69 | 518 | 33 | 388 | 33 |
|  | PT_Piara_395 | 109 | 75 | 592 | 1389 | 160 | 34 | 126 | 121 | 85 | 123 | 85 | 1032 | 174 | 9 | 69 | 518 | 33 | 388 | 33 |

**Table A3. Unified framework for quantifying abundance-based diversity facets (taxonomic, functional, and phylogenetic) using generalized Hill numbers (with the parameter *q* defining the sensitivity to entities abundances).** After Chao *et al*. (2014).

| **Facet of diversity** | **Element** | **Facet value** | **Entity** | **Relative abundance of an entity** | **Attribute (facet) diversity** |
| --- | --- | --- | --- | --- | --- |
| Taxonomic diversity | Species *i* = {1, …, S} | Unity for species *i* | Taxonomic entity (species identity) | *p_i_,* species relative abundance | ${{}^{q}{TD=}\left[ \sum_{i=1}^{S} 1\times\left( \frac{p_{i}}{\sum_{k=1}^{S} p_{k}} \right)^{q} \right]}^{\frac{1}{1-q}}$ |
| Functional diversity | Pairs of species (*i,j*) = {(*i,j*);*i,j* = 1, …, S} | Euclidean distance for pairs of species *(i,j)* | Functional entity (pairwise species unit of distance) | *p_i_ p_j_ /Q*, where *Q* = $\sum_{i,j=1}^{S} d_{ij}p_{i}p_{j}$ | ${{}^{q}{FD=}\left[ \sum_{i,j=1}^{S} d_{ij}\times\left( \frac{p_{i}p_{j}}{Q} \right)^{q} \right]}^{\frac{1}{1-q}}$ |
| Phylogenetic diversity | Tree branch *i* segment = {1, …, B} | Specie *i* tree branch length *L_i_* | Phylogenetic entity (branch unit of length) | $\frac{\boldsymbol{a}_{\boldsymbol{i}}}{\boldsymbol{T}}$**,** where $a_{i}$ = branch abundance, T= $\sum_{j=1}^{B} L_{j}a_{j}$ | ${{}^{q}{PD=}\left[ \sum_{i=1}^{B} L_{i}\times\left( \frac{a_{i}}{T} \right)^{q} \right]}^{\frac{1}{1-q}}$ |

**Table A4. Pairwise Pearson**′**s correlations between the three facets of diversity (taxonomic, functional, and phylogenetic) for different Hill numbers**′ **orders.** *: significant correlation (p<0.05); **: (p<0.01); ***: (p<0.001). Light grey cells indicate lack of functional data for Madidi and Pilón-Lajas transects.

| **Transect** | **Hill order** | **Taxonomic vs functional** | **Taxonomic vs phylogenetic** | **Functional vs phylogenetic** |
| --- | --- | --- | --- | --- |
| **Podocarpus** | *q* = 0 | 0,93 *** | 0,93 *** | 0,85 *** |
|  | *q* = 1 | 0,94 *** | 0,34 | 0,35 |
|  | *q* = 2 | 0,93 *** | -0,27 | -0,15 |
| **Río Abiseo** | *q* = 0 | 0,97 *** | 0,98 *** | 0,97 *** |
|  | *q* = 1 | 0,99 *** | 0,65 *** | 0,60 *** |
|  | *q* = 2 | 0,98 *** | 0,03 | -0,01 |
| **Madidi** | *q* = 0 |  | 0,95 *** |  |
|  | *q* = 1 |  | 0,81 *** |  |
|  | *q* = 2 |  | 0,55 ** |  |
| **Pilón-Lajas** | *q* = 0 |  | 0,96 *** |  |
|  | *q* = 1 |  | 0,48 * |  |
|  | *q* = 2 |  | -0,11 |  |
| **All sites** | *q* = 0 | 0,80 *** | 0,93 *** | 0,80 *** |
|  | *q* = 1 | 0,83 *** | 0,58 *** | 0,47 *** |
|  | *q* = 2 | 0,83 *** | 0,03 | -0,04 |

**Table A5. Model-averaged estimates, standard errors and p-values for selected variables in the best fitted models of taxonomic, functional, and phylogenetic diversity values calculated for Hill numbers of different order relative to distinct predictors for 114 plots of Andean TMFs.***: significant correlation (p<0.05), **: (p<0.01); ***: (p<0.001).

|  | **Hill order** | **Model parameters** | **Estimate** | **S.E.** | **p-val** |
| --- | --- | --- | --- | --- | --- |
| **Taxonomic diversity** | *q = 0* | Intercept | 4,80 | 0,08 | 2∙10^-16^ *** |
|  |  | Pilón-Lajas | 0,16 | 0.07 | 0.01 * |
|  |  | Podocarpus | 0.34 | 0.06 | 9.90∙10^-8^ *** |
|  |  | Río Abiseo | 0,22 | 0.06 | 7.87∙10^-4^ *** |
|  |  | Elevation | -4.31∙10^-4^ | 3.16∙10^-5^ | 2∙10^-16^ *** |
|  | *q* = 1 | Intercept | 4.68 | 0.08 | 2∙10^-16^ *** |
|  |  | Madidi : Elevation | -7.23∙10^-4^ | 4.92∙10^-5^ | 2∙10^-16^ *** |
|  |  | Pilón-Lajas : Elevation | -5.96∙10^-4^ | 4.70∙10^-5^ | 2∙10^-16^ *** |
|  |  | Podocarpus : Elevation | -4.57∙10^-4^ | 4.85∙10^-5^ | 2∙10^-16^ *** |
|  |  | Río Abiseo : Elevation | -5.57∙10^-4^ | 4.82∙10^-5^ | 2∙10^-16^ *** |
|  | *q* = 2 | Intercept | 4.29 | 0.11 | 2∙10^-16^ *** |
|  |  | Madidi : Elevation | -7.56∙10^-4^ | 6.06∙10^-5^ | 2∙10^-16^ *** |
|  |  | Pilón-Lajas : Elevation | -6.20∙10^-4^ | 5.78∙10^-5^ | 2∙10^-16^ *** |
|  |  | Podocarpus : Elevation | -4.73∙10^-4^ | 5.94∙10^-5^ | 1.85∙10^-15^ *** |
|  |  | Río Abiseo : Elevation | -5.91∙10^-4^ | 5.94∙10^-5^ | 2∙10^-16^ *** |
| **Functional diversity** | *q = 0* | Intercept | 10.30 | 0.25 | 2∙10^-16^ *** |
|  |  | Río Abiseo | 1.17 | 0.34 | 5.19∙10^-4^ *** |
|  |  | Elevation | -4.13∙10^-4^ | 1.23∙10^-4^ | 7.40∙10^-4^ *** |
|  |  | Río Abiseo : Elevation | -3.45∙10^-4^ | 1.62∙10^-4^ | 3.25∙10^-2^ * |
|  | *q* = 1 | Intercept | 10.12 | 0.23 | 2∙10^-16^ *** |
|  |  | Río Abiseo | 0.47 | 0.15 | 2.12∙10^-3^ ** |
|  |  | Elevation | -7.96∙10^-4^ | 1.03∙10^-4^ | 1.09∙10^-14^ *** |
|  | *q* = 2 | Intercept | 9.41 | 0.26 | 2∙10^-16^ *** |
|  |  | Río Abiseo | 0.47 | 0.18 | 7.89∙10^-3^ ** |
|  |  | Elevation | -8.13∙10^-4^ | 1.18∙10^-4^ | 6.46∙10^-12^ *** |
| **Phylogenetic diversity** | *q = 0* | Intercept | 8.91 | 0.11 | 2∙10^-16^ *** |
|  |  | Pilón-Lajas | 1.7∙10^-2^ | 0.15 | 0.91 |
|  |  | Podocarpus | -0.13 | 0.14 | 0.36 |
|  |  | Río Abiseo | 1.80∙10^-2^ | 0.13 | 0.89 |
|  |  | Elevation | -3.49∙10^-4^ | 4.82∙10^-5^ | 4.5∙10^-13^ *** |
|  |  | Pilón-Lajas : Elevation | 7.54∙10^-5^ | 6.67∙10^-5^ | 0.26 |
|  |  | Podocarpus : Elevation | 1.35∙10^-4^ | 6.41∙10^-5^ | 0.03 * |
|  |  | Río Abiseo : Elevation | 1.69∙10^-5^ | 6.04∙10^-5^ | 0.78 |
|  | *q* = 1 | Intercept | 7.06 | 4.03∙10^-2^ | 2∙10^-16^ *** |
|  |  | Madidi : Elevation | -1.32∙10^-4^ | 2.11∙10^-5^ | 3.26∙10^-10^ *** |
|  |  | Pilón-Lajas : Elevation | -5.52∙10^-5^ | 2.07∙10^-5^ | 7.56∙10^-3^ ** |
|  |  | Podocarpus : Elevation | -6.13∙10^-5^ | 2.21∙10^-5^ | 5.50∙10^-3^ ** |
|  |  | Río Abiseo : Elevation | -8.31∙10^-5^ | 2.15∙10^-5^ | 1.09∙10^-4^ *** |
|  | *q* = 2 | Intercept | 6.35 | 3.28∙10^-2^ | 2∙10^-16^ *** |
|  |  | Madidi : Elevation | -1.60∙10^-5^ | 1.71∙10^-5^ | 0.35 |
|  |  | Pilón-Lajas : Elevation | 3.90∙10^-5^ | 1.68∙10^-5^ | 0.02 * |
|  |  | Podocarpus : Elevation | 1.94∙10^-5^ | 1.79∙10^-5^ | 0.28 |
|  |  | Río Abiseo : Elevation | 2.21∙10^-5^ | 1.74∙10^-5^ | 0.20 |
